# The common symbiosis pathway controls plant root microbiomes in a host-specific manner

**DOI:** 10.64898/2026.05.08.723321

**Authors:** Anna Martyn, Ib Thorsgaard Jensen, Camilla Lind Salomonsen, Zuzana Blahovska, Ke Tao, Harm Dings, Þuríður Nótt Björgvinsdóttir, Bent Tolstrup Christensen, Giles Oldroyd, Rasmus Waagepetersen, Marianne Glasius, Simona Radutoiu

**Affiliations:** Department of Molecular Biology and Genetics, Aarhus University, Aarhus, Denmark; Max Planck Institute for Plant Breeding Research, Cologne, Germany; Department of Mathematical Sciences, Aalborg University, Aalborg, Denmark; Department of Chemistry, Aarhus University, Aarhus, Denmark; Department of Biology, University of Copenhagen, Copenhagen, Denmark; Laboratory of Molecular Biology, Wageningen University, Wageningen, Netherlands; Department of Agroecology, Aarhus University, Aarhus, Denmark; Crop Science Centre, University of Cambridge, Cambridge, United Kingdom; Donald Danforth Plant Science Center, St. Louis, MO 63132, Missouri, USA

**Keywords:** Plant-microbe interactions, root microbiome, symbiosis signalling, root exudates, host specificity, soil fertilisation

## Abstract

Crop nutrition depends on plant-microbe interactions, yet it remains unclear whether conserved genetic pathways impose universal rules on root microbiome assembly across plant hosts. Here, we show that the Common Symbiosis Signalling Pathway (CSSP), a conserved genetic module controlling endosymbiosis with arbuscular mycorrhizal fungi and nitrogen-fixing bacteria, regulates root microbiome assembly in a host-specific manner across contrasting fertilisation regimes. Using *Lotus japonicus* and *Hordeum vulgare*, we demonstrate that mutations in orthologous CSSP genes remodel root bacterial communities in both species, but with distinct taxonomic outcomes. In *Lotus*, CSSP disruption reduces rhizobial colonisation and promotes niche replacement by commensal taxa, whereas in *Hordeum*, the same mutations broadly restructure bacterial lineages without converging on *Lotus*-like responses. Root exudate profiling reveals host-specific metabolic differences, particularly in phenylpropanoid (flavonoids and coumarins) and gibberellin pathways, linking CSSP activity to chemically distinct rhizosphere environments that correlate with divergent microbiome assembly patterns across hosts. Moreover, root bacterial community composition accurately predicts plant nutritional status, highlighting tight coupling between host physiology and microbiome composition. Together, our results show that conserved symbiosis signalling regulates root microbiome assembly, while host-specific metabolic environments determine taxonomic outcomes. This extends CSSP function beyond canonical endosymbioses and positions symbiosis signalling as a general determinant of plant-microbiome interactions with implications for crop nutrition.

**Significance Statement:** Root microbiomes influence plant nutrition, yet how conserved host genetic pathways controlling interactions with intracellular symbionts shape root-associated microbiome assembly across divergent plant species remains unresolved. The Common Symbiosis Signalling Pathway (CSSP), conserved across most land plants, regulates root microbiome composition in both a legume and a cereal, but with distinct taxonomic outcomes. These effects correlate with CSSP-dependent differences in root exudate chemistry and host metabolic profiles. Together, our results show that conserved symbiosis signalling operates within host-specific metabolic contexts, providing a framework for understanding why disruption of the same genetic pathway can lead to divergent microbiome configurations across plant species.

## Introduction

The steady population increase and high demand for food production put immense pressure on agricultural production systems, raising major concerns about an upcoming global food crisis (1, 2). Crop productivity is primarily constrained by the availability of soil nutrients, which are essential for plant growth but often occur in forms inaccessible to plants (3–5). Agricultural practices rely heavily on both animal manure and mineral fertilisers to maintain yields; however, mineral fertilisers are costly and often inaccessible in developing countries, and excessive application of either input can have severe environmental consequences (4, 5). To sustainably enhance crop yields, it is crucial to understand how plants naturally optimise nutrient uptake and leverage this knowledge to improve crop productivity while minimising agriculture’s ecological footprint.

Plants interact with soil microorganisms and host unique microbial communities around their roots, collectively referred to as the ‘root microbiome’ (6–8). Extensive research has demonstrated that the root microbiome contributes substantially to plant health and nutrition (9). In some cases, plants establish highly specialised symbiotic relationships with microbes to enhance nutrient acquisition. Two of the best-studied intracellular symbioses are arbuscular mycorrhizal symbiosis (AMS) and symbiotic nitrogen fixation (SNF), which are well characterised at the molecular and genetic levels. In both associations, the plant host provides photosynthetically derived carbon to its microbial partner in exchange for mineral nutrients, primarily phosphorus in AMS and nitrogen in SNF (10). AMS is highly prevalent in the plant kingdom, occurring in the majority of land plant species (11, 12). Under phosphate starvation, plants exude strigolactones into the soil (12–15), which arbuscular mycorrhizal fungi (AMF) perceive and respond to by releasing Myc-factors (11, 16). These signals are recognised by specialised plant receptors (12), triggering calcium oscillations that activate downstream symbiosis signalling, enabling AMF to colonise the root cortex and form highly branched intracellular feeding structures known as ‘arbuscules’ (10, 11, 17, 18). These structures provide an extended surface area for nutrient exchange, where the plant supplies carbon sources to the fungus in return for phosphorus (12, 19). In contrast, SNF with nitrogen-fixing bacteria is far less widespread, occurring mostly in legumes, with a few non-legumes forming nodules with *Frankia*, and the tree *Parasponia* representing the only non-legume that nodulates with rhizobia (20, 21). Legumes release flavonoids (10, 22, 23) that are detected by nitrogen-fixing bacteria in the soil and activate bacterial production of Nod-factors (23, 24). These signals are perceived by the plant, triggering symbiosis signalling, as seen in AMS, but leading to the development of specialised root-derived organelles called ‘nodules’, inside which nitrogen-fixing rhizobia are endocytosed (10). Nitrogen fixation is then facilitated by host-supplied sugars and tightly controlled oxygen levels mediated by leghemoglobin (25–29). Together, SNF and AMS are essential strategies enabling plants to acquire nutrients critical for growth and reproduction. Both symbioses are controlled by a shared genetic pathway known as the ‘Common Symbiosis Signalling Pathway’ (CSSP), which is conserved across land plants (10, 11, 30–33). The CSSP is fundamental to both symbioses and operates not only in legumes, which facilitate both SNF and AMS, but also in cereals, which establish only AMS (33). A key CSSP gene is SYMRK, encoding a receptor-like kinase that acts as the first shared component of the pathway, interacting with Myc-and Nod-factor receptors and associating with regulatory proteins to trigger the release of calcium (Ca^2+^) signals (34–37). These Ca^2+^ signals are decoded by the nucleus-localised calcium- and calmodulin-dependent serine/threonine protein kinase CCaMK, which is considered the central regulator of symbiosis signal transduction (38–41). CCaMK acts as pacemaker, interpreting Ca^2+^ oscillations to activate endosymbiotic programs (42). CCaMK activates a complex transcriptional network involving genes essential for one or both symbioses (40, 43–45). Among these, the GRAS-domain transcription factors Nodulation Signalling Pathway1 (NSP1) and NSP2 are crucial. These proteins regulate nodule formation and bacterial infection during SNF and play a key role in strigolactone biosynthesis during early AMS (10, 46–49). In conclusion, the CSSP is the central regulatory network governing both SNF and AMS. A growing research field focuses on elucidating individual CSSP gene functions to enhance plant nitrogen and phosphorus uptake, with the goal of improving agricultural productivity (36, 50–53).

Recent studies have begun exploring whether individual CSSP genes may influence not only the establishment of symbioses but also broader microbial associations. Mutations in CSSP genes have been shown to alter bacterial and fungal communities in legumes, highlighting a potential role for these genes beyond symbiosis (54–58). Whether similar effects occur in non-leguminous plants, such as cereals, remains unclear. Cereals lack the ability to facilitate SNF and often suffer from nitrogen deficiencies, making it particularly interesting to investigate whether the CSSP also influences colonisation by other beneficial soil bacteria, such as free-living nitrogen-fixing isolates. Here, we tested the hypothesis that the CSSP shapes root-associated microbial communities in both a legume (*Lotus japonicus*) and a cereal (*Hordeum vulgare*). Using mutants impaired in orthologous CSSP genes (*symrk*, *ccamk*, *nsp1*, *nsp2*), we examined how disrupted symbiosis signalling influences root microbiota composition, root exudate profiles, and plant growth. We combined these genetic perturbations with soils subjected to long-term differential nitrogen and phosphorus fertilisation. This revealed that soil nutrient status strongly affects root-associated bacterial communities and that microbiota composition can accurately predict the plant’s nutritional state. Independent of nutrient availability, active symbiosis signalling exerts a strong influence on bacterial community assembly in both hosts. In *Lotus*, CSSP mutants primarily lose symbiont colonisation, creating opportunities for other microbes to occupy available niches, whereas in *Hordeum*, colonisation by diverse bacterial taxa is broadly altered. These community shifts appear to be linked to CSSP-dependent changes in root exudation, particularly affecting phenylpropanoids. Together, our findings demonstrate that CSSP genes play a broader role in plant-microbe interactions beyond their known functions in SNF and AMS. These insights have implications for enhancing nutrient uptake and for the development of host-tailored microbiome management strategies.

## Results

### Long-term effects of soil fertilisation on soil bacterial communities

The soils used in this study were sampled from the Askov long-term field experiment in Denmark, where plots had been consistently treated with specific fertiliser combinations for over 125 years (59). We focused on three soils (hereafter soil types) that had been subjected to different treatments: (1) nitrogen, phosphorus, and potassium fertilisation (NPK); (2) nitrogen-deficient soil (PK); and (3) unfertilised soil (UF). All soils followed the same crop rotation and agricultural practices over time. Given the long-term fertilisation history, we first asked whether these treatments had influenced soil bacterial communities. Profiling the soil microbiota using 16S V5-V7 rRNA amplicon sequencing (Fig. 1) revealed similar within-sample diversity (alpha diversity, Chao1 index) across the three soils (Fig. 1B), but substantial differences in community composition (between-sample diversity, beta diversity; Bray-Curtis dissimilarities, Fig. 1C). Several well-known bacterial orders (60) were among the most abundant taxa (Fig. 1D), with some showing differential abundance across soil types; for example, *Pseudomonadales* were enriched in UF, whereas *Bacillales*, *Gaiellales*, and *Xanthomonadales* were more abundant in NPK soil (Fig. 1E). Similar numbers of amplicon sequence variants (ASVs; ∼1200) were identified in all soils. Of these, fewer than 40% were shared across soil types (Fig. 1F, Table S1). Both unique and shared ASVs spanned a broad range of bacterial orders, indicating that long-term fertilisation influenced community composition at both low and higher taxonomic levels (Fig. 1F). This demonstrates that long-term fertilisation maintains overall diversity while shaping bacterial community composition, resulting in distinct communities likely adapted to the specific soil conditions imposed by different fertilisation regimes.

**Figure 1.**
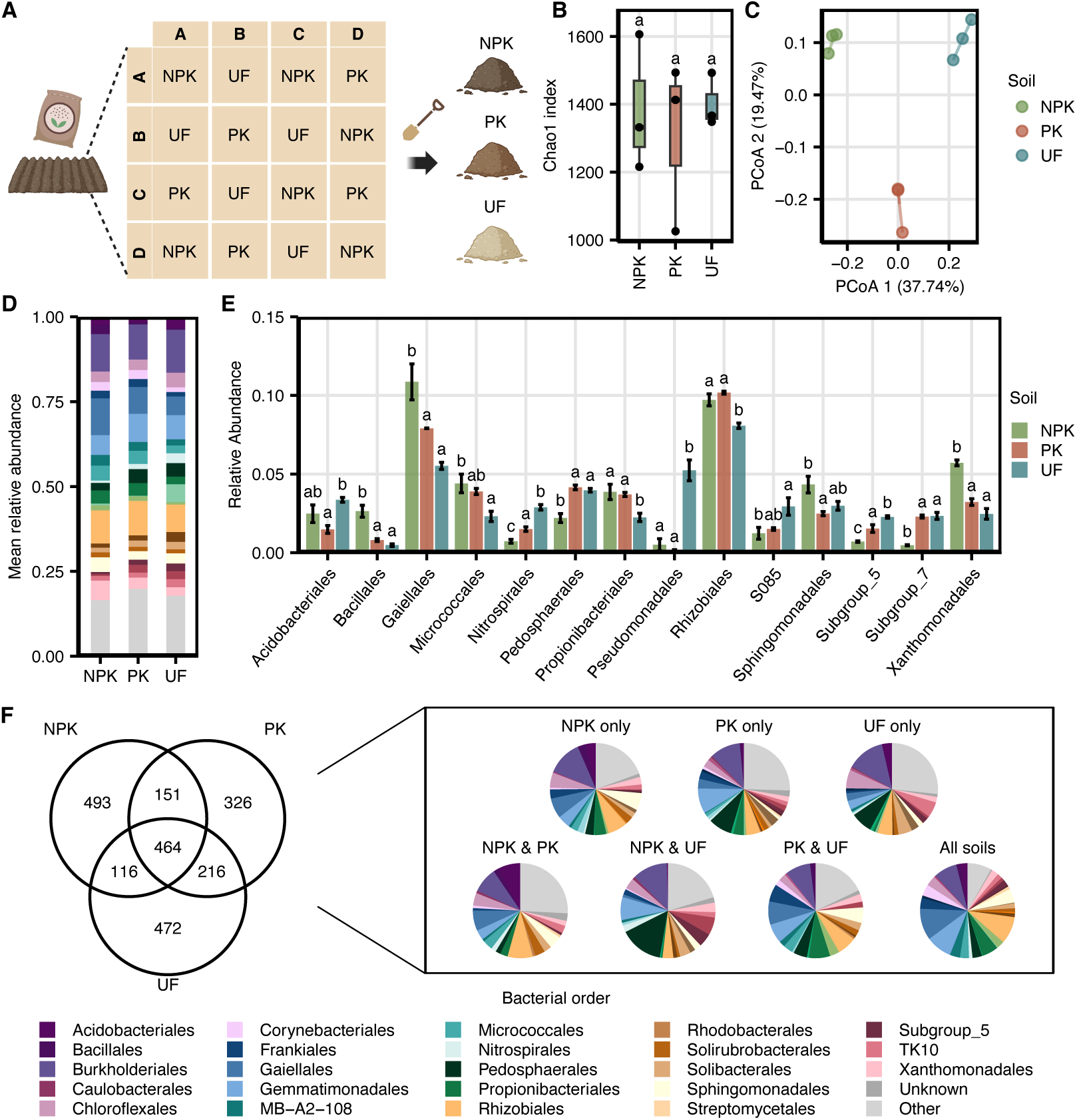
Long-time fertilisation shapes soil bacterial community composition. (A) Three soils were collected after 125 years of field fertilisation at the Askov long-term experiment (Christensen *et al*., 2022), including nitrogen, phosphorus, and potassium fertilised (NPK), phosphorus and potassium fertilised (PK), and unfertilised (UF) soil. Soils were subjected to 16S rRNA V5-V7 amplicon sequencing (*N* = 3 per soil). (B) Alpha diversity (Chao1 index) of bacterial communities across soils, calculated from a rarefied ASV table (minimum 1000 reads per sample). (C) Principal coordinates analysis (PCoA) based on Bray-Curtis distances. (D) Stacked bar plots showing the mean relative abundance of bacterial orders among the 20 most abundant orders in at least one soil type (remaining taxa grouped as “Other”). (E) Relative abundance of bacterial orders showing significant differences across soils. (F) Venn diagram showing overlap of ASVs detected across soils (present in ≥2 of 3 samples). Pie charts indicate taxonomic composition at order level for each section of the Venn diagram. Statistical significance was assessed using one-way ANOVA followed by Tukey’s HSD post hoc test (P < 0.05).

### Host-specific colonisation by distinct soil microbiota members

Next, we cultivated *Lotus* and *Hordeum* wild-type in the three soils (NPK, PK and UF) for three weeks (Fig. 2A) and assessed their phenotype and associated microbiota. Shoot fresh weight revealed that, as expected (61), fertilisation levels had a significant effect on plant growth, with reduced growth as nutrient availability decreased (Fig. 2B). *Lotus* formed pink root nodules in all soils, although nodule numbers were significantly higher in PK and UF than in NPK, reflecting nitrogen repression of symbiosis (Fig. 2C). The highest nodule counts were observed in PK, likely due to sufficient phosphorus supporting SNF (62). We collected rhizosphere, root, and nodule samples for microbiome analysis. As expected, alpha diversity was highest in rhizosphere samples of both hosts, followed by root and then nodule samples (Chao1 index, Fig. S1A,B) (55). Fertilisation history did not affect alpha diversity in *Hordeum*, but *Lotus* rhizosphere communities exhibited reduced diversity in PK and UF soils (Fig. S1A,B), coinciding with higher nodule numbers (Fig. 2C), consistent with SNF contributing to rhizosphere microbiota structuring (54). Beta diversity was significantly influenced by both sample fraction and soil type in both plants (Fig. 2D), with clear separation of rhizosphere and root communities. *Lotus* nodule communities displayed similar beta diversity patterns across soils (Fig. 2D). Community structure analysis revealed common features in both hosts. *Burkholderiales* was the most abundant order, while *Rhizobiales* was particularly enriched in *Lotus* root and nodule fractions due to highly competitive symbionts (Fig. 2E, Fig. S1C,D). Additional host-specific colonisation patterns were observed, including a higher abundance of *Pseudomonadales* in *Lotus*, whereas *Flavobacteriales* and *Streptomycetales* were more prevalent in *Hordeum*, with the latter particularly enriched in roots. Soil-dependent community shifts were also detected. For instance, *Burkholderiales* were the most abundant in PK and UF in both hosts. In *Lotus*, *Chloroflexales* were enriched in rhizosphere and root fractions when plants were grown in UF soil, while *Streptomycetales* (rhizosphere) were more abundant in NPK. In *Hordeum*, *Caulobacterales* were enriched in NPK roots, while *Flavobacteriales* increased in PK and UF (Fig. 2E, Fig. S1C). *Lotus* nodules were colonised by *Mesorhizobium* and *Pseudomonas* ASVs, which largely overlapped between soils but differed in individual abundances in the nodules (Fig. S1D). Together, these results show that long-term fertilisation regimes significantly impact plant fitness and associated root microbiota, and that different hosts adapt to the given differences and selectively recruit specific bacterial members, reinforcing the concept of host preference (63).

**Figure 2.**
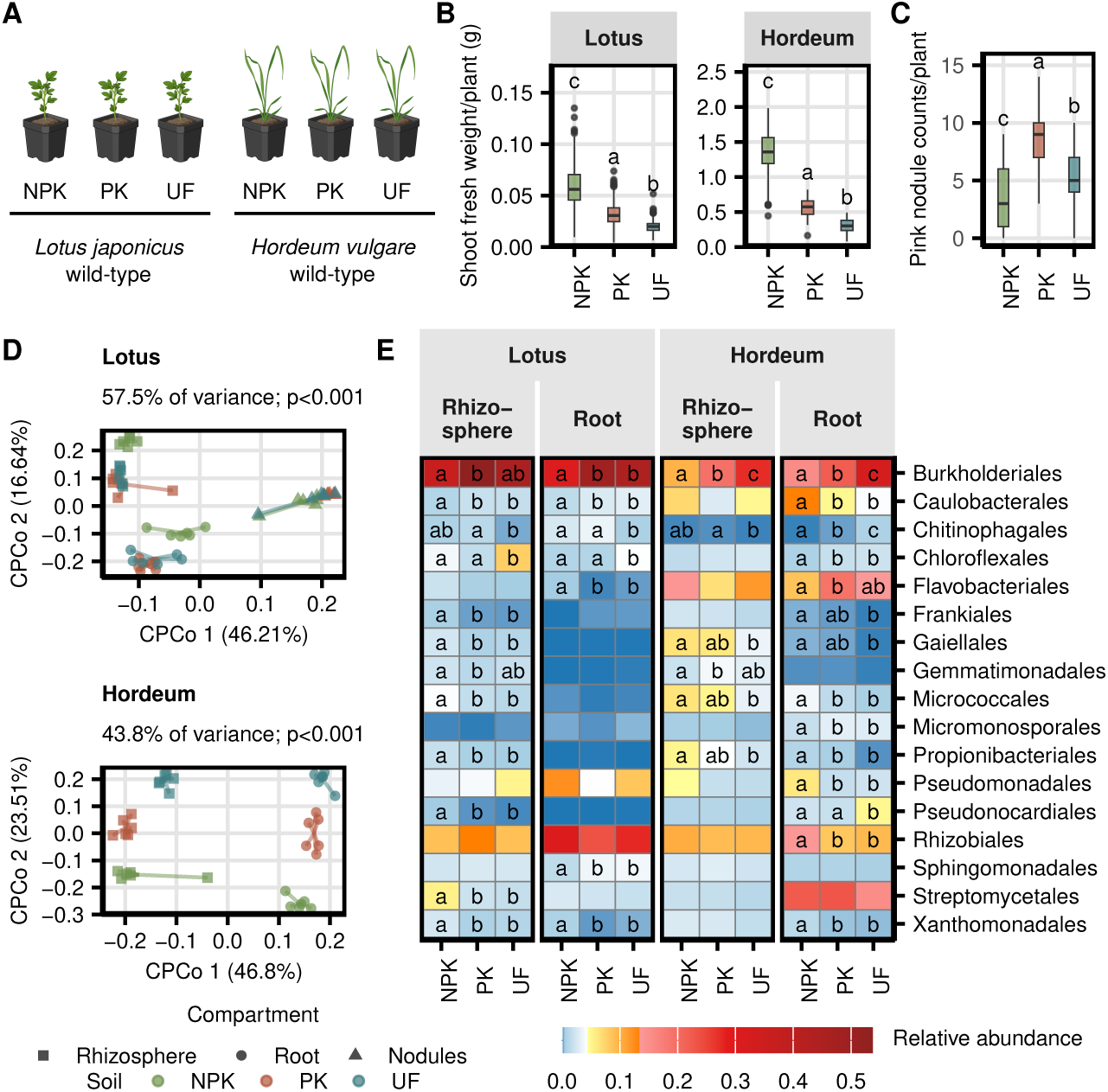
Soil legacy shapes plant growth and root-associated bacterial communities in *Lotus japonicus* and *Hordeum vulgare*. (A) Wild-type *Lotus* and *Hordeum* were grown for three weeks on NPK, PK, or UF soils. Each replicate consisted of three pots containing 10 plants per pot for *Lotus* (*N* = 60) and one plant per pot for *Hordeum* (total *N* = 18). Plants were pooled per replicate to collect rhizosphere, root, and nodule compartments for microbiome profiling using 16S rRNA V5-V7 amplicon sequencing (*N* = 6 per plant-compartment combination). (B) Shoot fresh weight (g) per plant. (C) Number of pink nodules per *Lotus* plant. (D) Constrained principal coordinates analysis (CPCoA) of Bray-Curtis distances showing bacterial community composition in rhizosphere, root, and nodule (for *Lotus*) compartments. Shapes indicate compartment, and colours indicate soil. (E) Heatmap of the mean relative abundance (RA) of orders with mean RA ≥1% across all rhizosphere and root samples for *Lotus* or *Hordeum*. Lower-abundance orders are grouped as “Other”. Letters indicate significant differences between soils within each plant-compartment (one-way ANOVA, Tukey’s HSD, P < 0.05).

### High accuracy prediction of plant nutrient status based on bacterial community composition

Following the observed large differences in rhizosphere and root microbiota across the different soils, we asked whether these microbial signatures could be used to predict the nutritional state of *Lotus* and *Hordeum*. We developed a two-step classification model trained on bacterial community profiles from plants grown in NPK, PK, and UF soils. The first model distinguishes plants grown in NPK soil from those in non-NPK soil, while the second model further separated PK from UF samples (Fig. 3A). Predictions were based on log-ratios of relative abundances of specific ASV combinations from rhizosphere or root microbiomes (Fig. 3C; Table S2), which together accounted for 4-36% of total relative abundance (Fig. 3D). Remarkably, the model achieved an overall accuracy of 93.3% using *Lotus* and 80-83.3% using *Hordeum* rhizosphere and root microbiome data, with only 15 out of 120 samples not previously seen by the model being misclassified (Fig. 3B,C). For *Lotus*, influential ASVs belonged to *Burkholderiales* (*Oxalobacteraceae*, *Comamonadaceae*), along with *Pseudomonadales* and *Rhizobiales*, each displaying compartment- and soil-type-specific associations (Fig. 3D, Table S2). In *Hordeum*, a broader taxonomic range contributed to prediction accuracy, with *Burkholderiales*, *Caulobacterales*, *Flavobacteriales*, *Pseudomonadales*, and *Streptomycetales* ASVs particularly important across all soil-compartment combinations. *Bacillales* and *Cellvibrionales* ASVs were additional predictors of NPK and PK cultivation, respectively, in rhizosphere samples, while *Pseudonocardiales* ASVs were used for PK predictions for root samples (Fig. 3D, Table S2). Predictive ASVs differed substantially between the two hosts and were only partially conserved across fertilisation regimes, suggesting that each plant species recruits distinct microbial indicators of nutrient status rather than relying on universal predictors. In summary, our model based on bacterial community profiles provides a robust tool for predicting plant nutrient status with high accuracy, highlighting the plant-specific recruitment of indicator taxa and the intricate interplay between host species, soil type, and microbial community composition.

**Figure 3.**
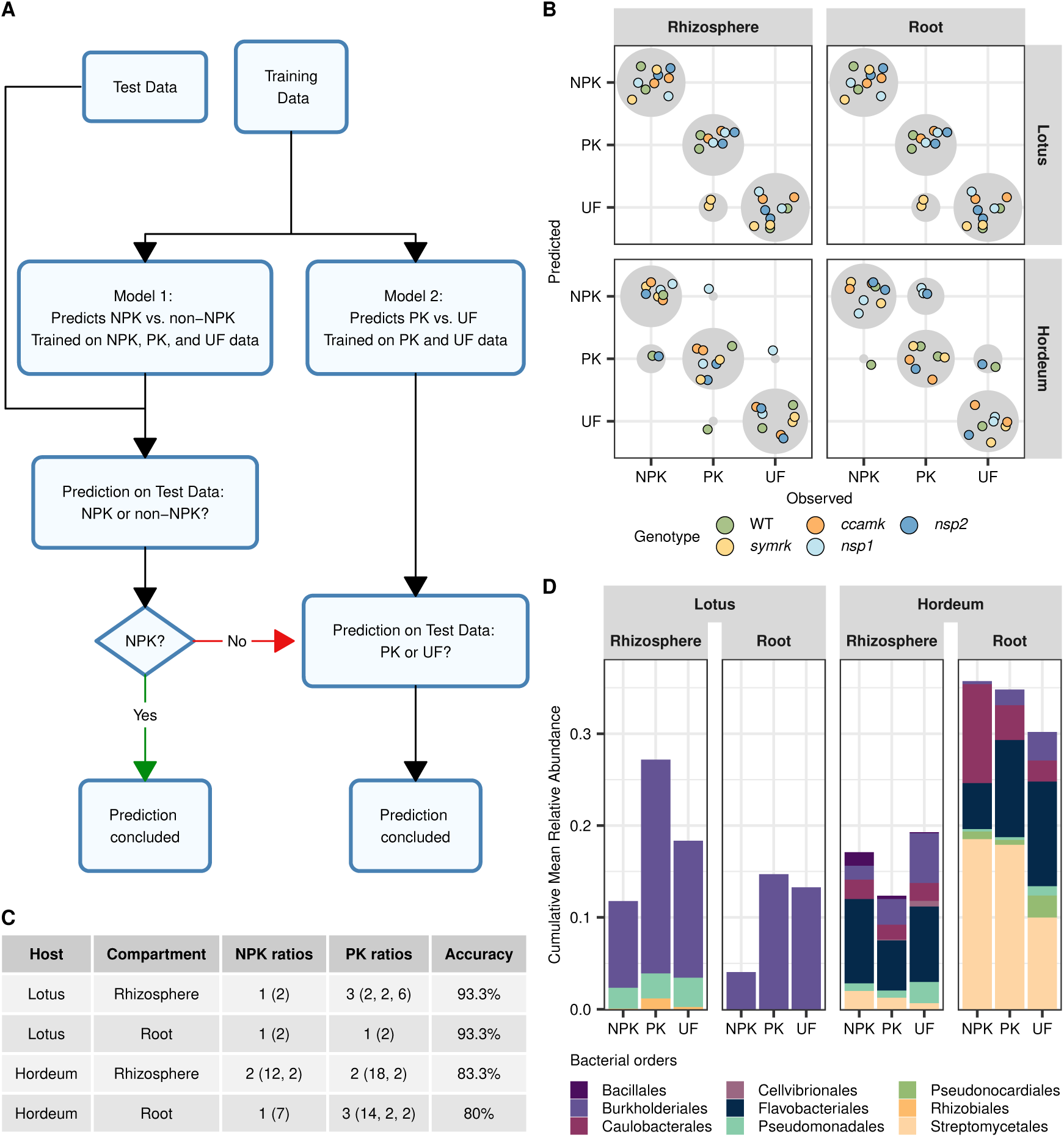
Prediction of plant nutrient status from bacterial community composition. (A) Schematic of the nested modelling approach used to predict soil type based on bacterial ASV profiles. Two binary classifiers were fitted using codacore: the first distinguishes NPK from non-NPK soils, and the second distinguishes PK from UF soils. Separate models were trained for *Lotus japonicus* and *Hordeum vulgare*, and for rhizosphere and root fractions. Two-thirds of samples were used for training and one-third for testing with balanced representation of soils and genotypes (wild-type, *symrk*, *ccamk*, *nsp1*, *nsp2*). (B) Prediction results on the test dataset. The x-axis indicates true soil type and the y-axis indicates predicted soil type. Each point represents one sample; bubbles on the diagonal indicate correct predictions and off-diagonal bubbles indicate misclassifications. (C) Log-ratio features selected by each model, showing the number of ASVs included in each log ratio (numbers in parantheses) and prediction accuracy for the test dataset. Model 1 corresponds to NPK *vs*. non-NPK classification (NPK ratios) and Model 2 corresponds to PK *vs*. UF classification (PK ratios). (D) Mean cumulative relative abundance of ASVs involved in prediction (all log-ratio components across both models) grouped by bacterial order.

### Impairment of symbiosis signalling genes alters microbiome assemblies

Building on the observed effects of soil nutrient levels and host-determined root-associated communities, we tested whether active symbiosis signalling in wild-type *Lotus* and *Hordeum* contributes to the assembly of their associated microbial communities. To address this, we compared CSSP mutants (*symrk*, *ccamk*, *nsp1*, and *nsp2*) cultivated in the three soil types with their corresponding wild-type plants (Fig. 4A). In *Lotus*, *ccamk* and *nsp1* mutants showed significantly reduced growth in NPK soil, while all mutants exhibited reduced growth in PK and UF soils (Fig. 4B). This growth reduction likely reflects the loss of SNF, consistent with their defective symbioses (Fig. S2A) (46, 56, 64–67, 10). In *Hordeum*, growth was significantly reduced in *ccamk* mutants across all soils, whereas other mutants showed milder effects (Fig. 4B). This reduction was not observed under axenic conditions (Fig. S2B), indicating that the phenotype depends on the presence of soil microbes. Alpha diversity was generally unaffected by CSSP impairment in *Lotus*, whereas in *Hordeum* it was reduced in the rhizosphere of most mutants (NPK soil) and in *ccamk* roots (UF soil) (Fig. S3A,B). Beta diversity was primarily driven by soil type (Fig. 4C, Fig. S3C,D), although differential clustering of genotypes was observed across plant-soil-fraction combinations (Fig. 4C, Fig. S3E,F). Analysis of cumulative relative abundances at the bacterial order level revealed genotype- and soil-specific patterns across wild-type and CSSP mutants (Fig. S4A,B). In the *Lotus* rhizosphere, symbiosis impairment affected the abundance of several bacterial orders, while in roots, the order *Rhizobiales* in particular was reduced in all mutants across all soils, likely due to impaired symbiont colonisation (Fig. S4B). In *Hordeum*, the strongest effects occurred in *ccamk* and *nsp* mutants in both rhizosphere and root fractions, involving taxa such as *Burkholderiales*, *Caulobacterales*, *Pseudomonadales*, and *Rhizobiales* (Fig. S4B).

**Figure 4.**
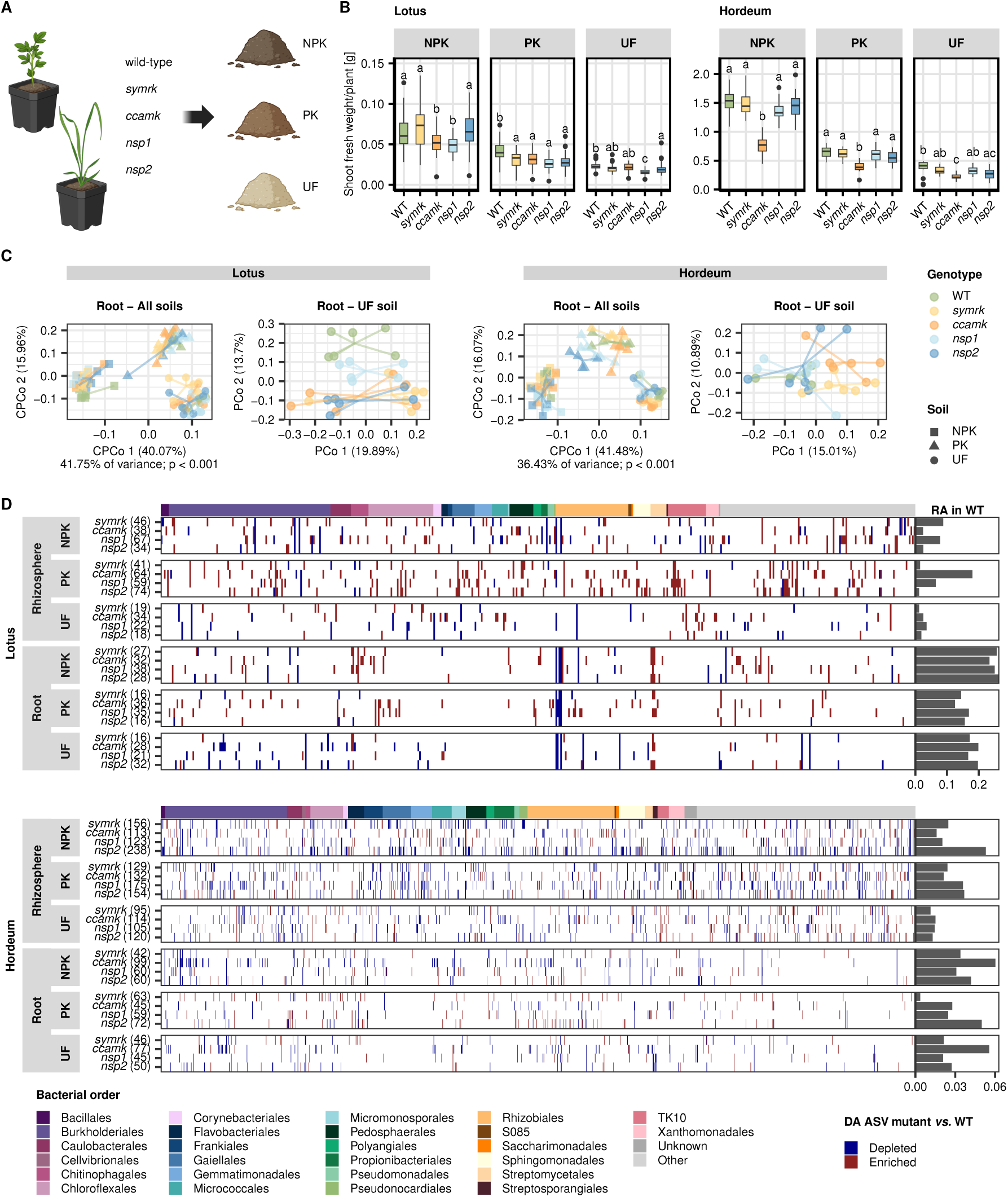
Symbiosis signalling shapes host-specific plant growth and root-associated microbiota. (A) *Lotus* and *Hordeum* wild-type and symbiosis mutants (*symrk*, *ccamk*, *nsp1*, *nsp2*) were grown for three weeks in NPK, PK, or UF soils. Rhizosphere and root microbiome samples were collected as described in Figure 2. (B) Shoot fresh weight (g) per plant (*N* = 60 for *Lotus* and *N* = 18 for *Hordeum* across most genotypes). (C) Constrained PCoA of Bray-Curtis distances showing bacterial community composition in root samples across all soils, with colours indicating genotype and shapes indicating soil type (left), and unconstrained PCoA of root samples from UF soil coloured by genotype (right). Extended PCoA plots, including rhizosphere and individual soil compartments, are shown in Fig. S3C-F. (D) Heatmap of ASVs with differential abundance (DA ASVs; identified via MaAsLin2 and structural zero analysis) in at least one mutant relative to wild-type across compartment-soil combinations. Depleted ASVs are shown in blue, enriched ASVs in red. The top bar indicates taxonomic assignment at the order level (bacterial orders among the 20 most abundant in *Lotus* or *Hordeum* shown; others grouped as “Other”). Numbers in brackets indicate the total DA ASVs per compartment-soil-genotype combination. Bars to the right show the mean cumulative relative abundance of DA ASVs in the corresponding wild-type compartment.

Differential abundance analysis (Fig. 4D, Table S3) identified ASVs representing up to approximately 6-27% of the total relative abundance in wild-type roots that showed altered colonisation in CSSP mutants in both hosts. These ASVs spanned a broad taxonomic range. In *Lotus*, all mutants displayed reduced colonisation of *Rhizobiales* ASVs, most likely symbiotic rhizobia, alongside other taxa that showed genotype-specific shifts. In *Hordeum*, several *Burkholderiales*, *Micromonosporales*, *Propionibacteriales*, and *Rhizobiales* ASVs were consistently reduced in all mutants (or at least in *ccamk* and *nsps*), while other taxa showed mutant-specific enrichment or depletion patterns.

Overall, impairment of the CSSP at different stages not only impacted plant growth but also significantly influenced the colonisation of a diverse range of bacteria in the *Lotus* and *Hordeum* rhizosphere and roots. The distinct microbiome shifts observed across individual mutants indicate that CSSP genes contribute more broadly to plant-microbe interactions beyond their established symbiotic roles. The particularly strong growth defect of *Hordeum ccamk* in the presence of microbes suggests an additional, yet poorly characterised, function of this gene in regulating plant performance when grown in complex microbial environments.

### Symbiosis signalling pathways can be linked to differential root exudation in both hosts

Root exudates are key drivers of interactions at the root-soil interface, forming a diverse chemical pool that fosters microbial competition and cooperation (68–70). Beyond serving as a nutrient source, root exudates contain defence-related metabolites with antimicrobial properties, as well as signalling molecules that are important for communication with beneficial microbes (9, 70). Previous work has shown that NSP1/NSP2 genes modulate strigolactone production, and CCaMK modulates phenylpropanoid biosynthesis during symbiosis (42, 48). To assess whether the observed effects of CSSP mutations on root-associated bacterial communities in *Lotus* and *Hordeum* are linked to alterations in root exudates, we analysed root exudate composition in CSSP mutants and wild-type plants under sterile, low-nutrient conditions using non-targeted liquid chromatography-mass spectrometry (LC-MS/MS). Due to differences in plant phenology, two distinct methods for extracting root exudates were used for *Lotus* and *Hordeum*; hence, only comparisons between genotypes of the same species are presented here.

After removal of background signals based on negative controls, we detected 661 and 971 metabolic features across all genotypes in *Lotus* and *Hordeum*, respectively (Tables S4,5). These metabolites spanned diverse chemical classes, including alkaloids, amino acids and peptides, carbohydrates, fatty acids, polyketides, shikimates and phenylpropanoids, as well as terpenoids (Fig. S5). The exudate profiles of all CSSP mutants, except for *Hordeum symrk*, displayed distinct clustering relative to their respective wild-type (Fig. 5A).

**Figure 5.**
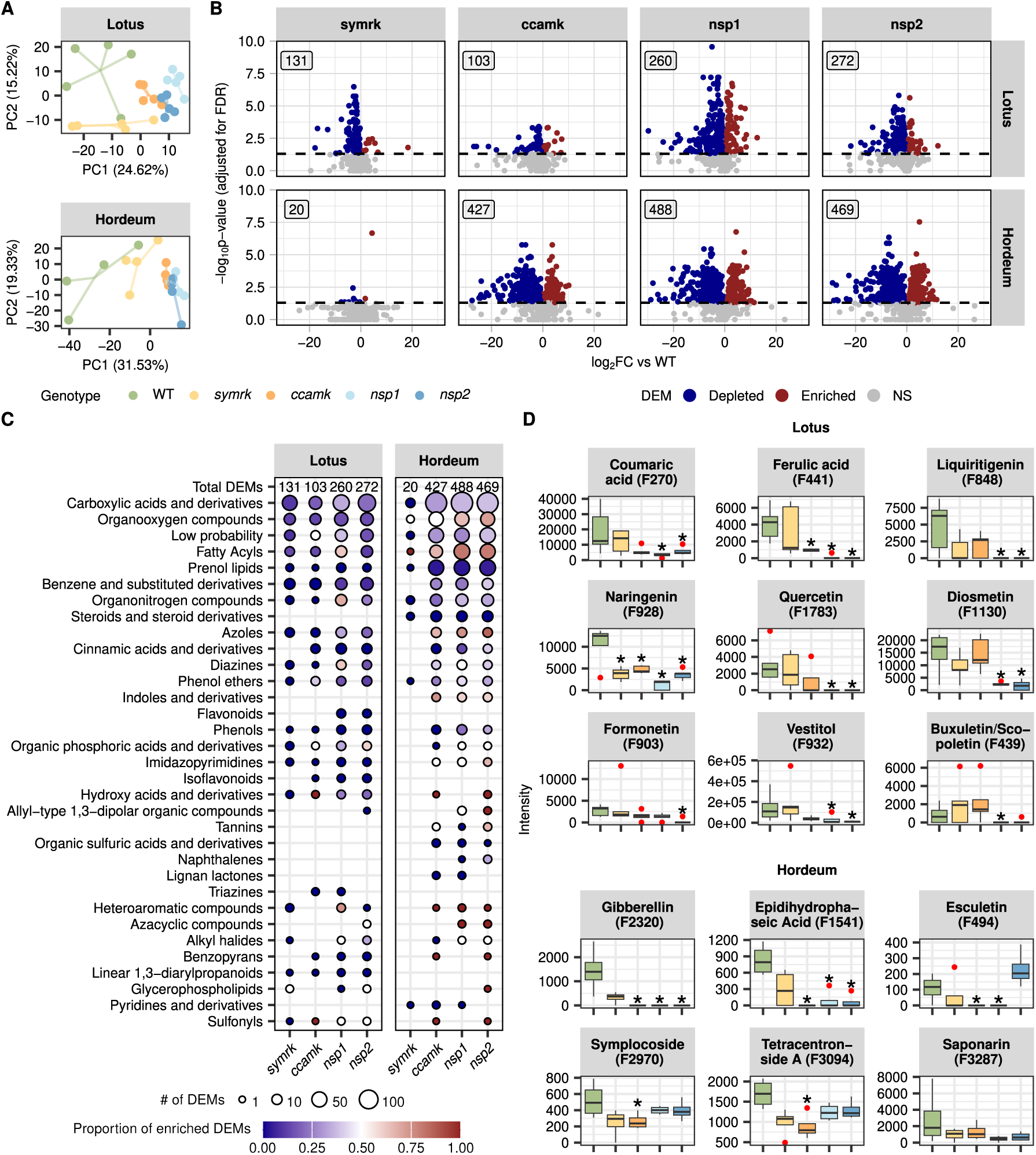
Root exudation profiles are altered in CSSP mutants of *Lotus* and *Hordeum*. *Lotus* and *Hordeum* wild-type and CSSP mutants were cultivated under microbe-free conditions (*Lotus*: *N* = 6, 12 plants per replicate; *Hordeum*: *N* = 6, one plant per replicate) and root exudates were collected for non-targeted LC-MS/MS analysis. (A) Principal component analysis (PCA) of all detected metabolites (background and low-frequency features removed), separated by host species, with colours indicating genotypes. (B) Volcano plots showing differential exudation of metabolites for each mutant relative to wild-type, with x-axis representing log_2_ fold change and y-axis showing −log_10_ p-value adjusted for false discovery rate (FDR) with Benjamini Hochberg. Dots are coloured by depletion (blue), enrichment (red), or not significant (gray). The number in each panel indicates the total number of significantly differentially exuded metabolites (DEMs) for the genotype. Features with estimated log_2_ fold-change above 30 were omitted for visual clarity. (C) Bubble plot summarising the total number of DEMs per genotype (x-axis) and their distribution across chemical classes (ClassyFire annotation). Bubble size reflects the number of DEMs per class, and bubble colour indicates the proportion of mostly depleted (blue), mixed (white), or mostly enriched (red) metabolites relative to wild-type. Classes where at least two metabolites are differentially exuded in at least one mutant compared to wild-type are shown. (D) Boxplots for selected DEMs with high-confidence structure annotations, showing intensity across genotypes; asterisks indicate significant differences relative to wild-type (Tobit regression with likelihood ratio tests, P < 0.05 after adjusting for multiple testing with Benjamini Hochberg).

In *Lotus*, shikimates and phenylpropanoids were significantly reduced in CSSP mutants, particularly in *nsps* compared to wild-type (Fig. S5). We identified between 103 and 272 metabolites with differential intensity in individual *Lotus* CSSP mutants relative to the wild-type (hereafter differentially exuded metabolites, DEM), spanning multiple compound classes and indicating a broad impact of CSSP mutations (Fig. 5B,C, Table S6). Notably, phenylpropanoids, including cinnamic acids and derivatives, isoflavonoids, and flavonoids, were strongly reduced across mutants (Fig. 5C). Structural predictions identified several well-known components of the phenylpropanoid pathway (71, 72), including coumaric acid (Feature270), ferulic acid (441), liquiritigenin (848), naringenin (928), quercetin (1783), diosmetin (1130), formonetin (903), vestitol (932), and buxuletin (also known as scopoletin) (439), most of which were present at lower levels in *ccamk* and *nsp* mutants compared to wild-type plants (Fig. 5D).

In *Hordeum*, the wild-type and *symrk* exudate profiles were similar, whereas *ccamk* and *nsp* mutants exhibited pronounced metabolic shifts (Fig. 5A). The number of DEM was low in *symrk* mutants (20), but up to 488 DEM were detected in other mutants (Fig. 5B), spanning several compound classes (Fig. 5C, Table S7). Well-characterised compounds included gibberellin (2320), the abscisic acid catabolite epidihydrophaseic acid (1541), and several phenylpropanoids: the coumarin esculetin (494), and the flavonoids symplocoside (2970), tetracentronside A (3094), and saponarin (3287) (Fig. 5D). Most of these compounds were less abundant in *ccamk* and/or *nsp* mutants compared to wild-type plants; however, this difference was not statistically significant for saponarin.

Although direct comparisons between species cannot be made due to differences in plant phenology, phenylpropanoids were consistently affected in CSSP mutants of both *Lotus* and *Hordeum* (Fig. 5C,D), suggesting a conserved role of CSSP genes in regulating their biosynthesis. However, the specific phenylpropanoid compounds altered were host-specific. In *Lotus*, isoflavonoid biosynthesis was primarily affected, whereas in *Hordeum*, coumarins and glycosylated flavonoids showed the strongest changes. Beyond phenylpropanoids, certain metabolites such as the plant hormone gibberellin were modulated in *Hordeum*, whereas these compounds did not show significant changes in *Lotus* (considering only metabolites with confidently assigned molecular formulas).

In summary, CSSP mutations affect hundreds of metabolites across diverse chemical classes. Some shifts are conserved across hosts (phenylpropanoids), whereas others are species-specific (isoflavonoids in *Lotus*; gibberellins in *Hordeum*) and genotype-specific. These findings suggest that CSSP genes regulate not only the biosynthesis of well-characterised symbiosis-related compounds but also broader metabolic processes. The observed genotype- and species-specific differences in exudate profiles could explain the shifts in root-associated microbial communities.

### Community shifts were replicated in gnotobiotic systems with synthetic bacterial communities

Our findings demonstrate a significant effect of symbiosis signalling on bacterial community establishment in *Lotus* and *Hordeum* grown in soil, as well as on their root exudate profiles under sterile conditions. However, soil is a complex environment, making it difficult to dissect individual plant-microbe interactions and uncover possible mechanisms underlying the observed community shifts. We therefore asked whether these effects could be recapitulated under controlled experimental conditions using complex but defined bacterial communities. For this, we employed controlled growth systems (Materials and methods) with reconstituted synthetic communities (SynComs) to determine whether the effects of impaired symbiosis signalling on bacterial colonisation in *Lotus* and *Hordeum* are directly plant-mediated, likely via altered root exudation, and independent of other soil environmental factors. We used SynComs composed of bacterial isolates derived from healthy, soil-grown *Lotus* and *Hordeum* plants, representing native members of their respective core microbiota (*Lotus* collection: Lj-SPHERE (63); *Hordeum* collection: (73); Methods; Tables S8, 9). Wild-type plants and CSSP mutants (*symrk*, *ccamk*, *nsp1*, *nsp2*) were cultivated under sterile, low-nutrient conditions and inoculated with either the Lj-SPHERE SynCom (291 isolates) or the *Hordeum* SynCom (316 isolates), respectively (Fig. 6A). Rhizosphere and root communities were analysed after three weeks using 16S rRNA V5-V7 amplicon sequencing, alongside plant phenotypic measurements.

**Figure 6.**
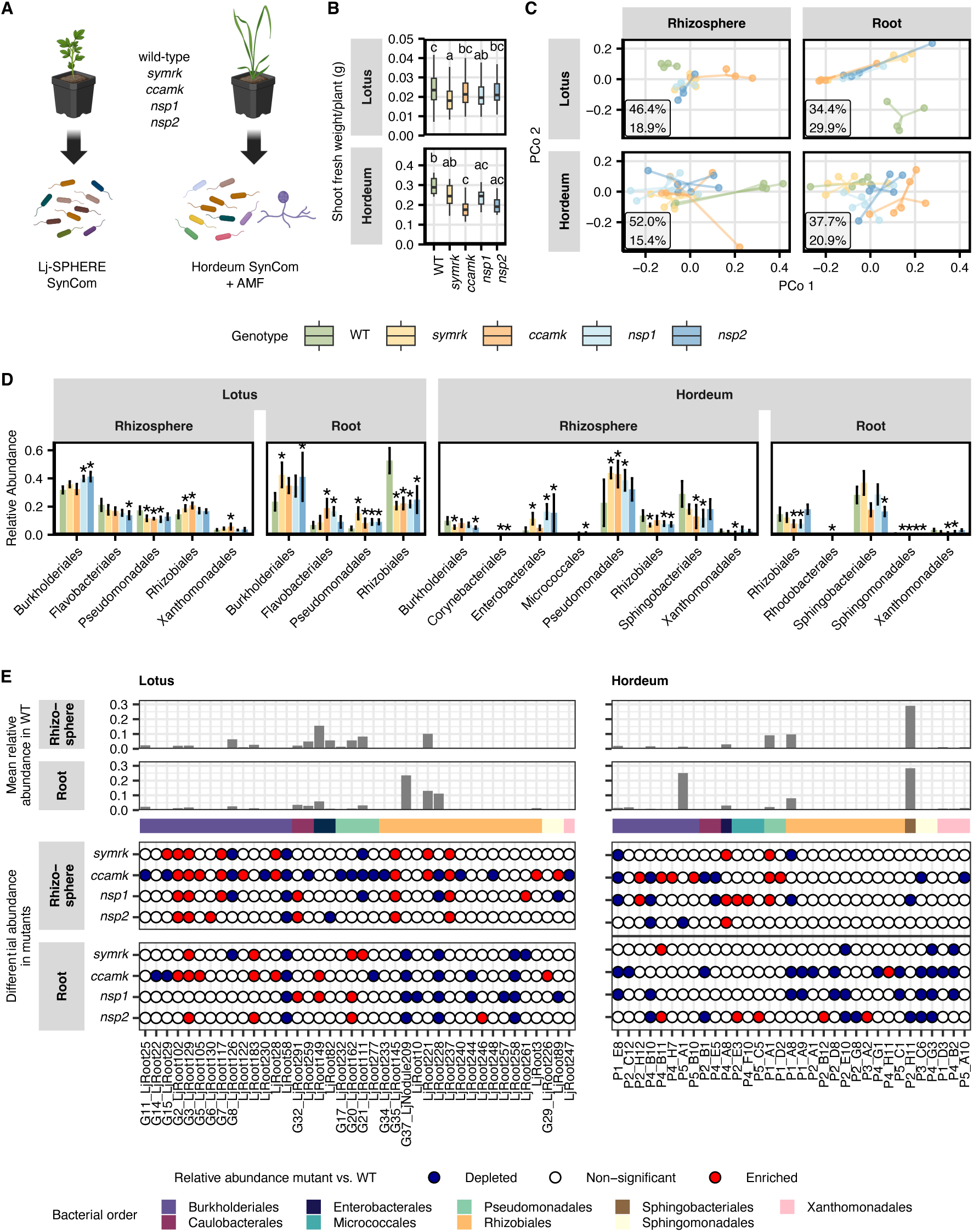
Synthetic community reconstitution experiments reveal plant-mediated shifts in root microbiota of symbiosis mutants. (A) *Lotus* and *Hordeum* wild-type and symbiosis mutants (*symrk*, *ccamk*, *nsp1*, *nsp2*) were grown for three weeks in gnotobiotic systems supplemented with *Lotus*- or *Hordeum* derived bacterial culture collections, with AMF additionally applied in the *Hordeum* approach (Methods, Tables S8,S9). Rhizosphere and root samples were collected and subjected to 16S rRNA V5-V7 amplicon sequencing (*Lotus*: *N* = 4, 25 seedlings per biological replicate; *Hordeum*: *N* = 6, three plants per biological replicate). (B) Shoot fresh weight (g) per plant. (C) PCoA analysis based on Bray-Curtis distances showing bacterial community composition in rhizosphere and root samples across genotypes (colours). Boxes indicate the percentage of total variance explained by the first and second principal coordinates for each compartment. (D) Barplots showing cumulative relative abundance of bacterial orders that were significant altered in at least one mutant compared to wild-type, as determined by MaAsLin2 (FDR < 0.05) across compartments. (E) Differentially abundant isolates detected in at least one mutant relative to wild-type based on MaAsLin2 and structural zero analysis. Top panels show mean relative abundance in wild-type rhizosphere and root samples. The annotation bar indicates taxonomic assignment at the order level. Bubble plots indicate significant depletion (blue) or enrichment (red) of isolates in mutants relative to wild-type. Unmatched ASVs were excluded from the analysis.

In *Lotus*, all CSSP mutants exhibited reduced growth, with the strongest effects observed in *symrk* and *nsp1* mutants (Fig. 6B), consistent with results obtained in depleted soil conditions (Fig. 4B). None of the mutants developed root nodules, and thus, SNF formation was exclusive to *Lotus* wild-type (Fig. S6C). Alpha diversity of bacterial communities was generally comparable between the rhizosphere and root fractions, whereas nodule diversity was reduced, consistent with previous findings (Fig. S6A). Between *Lotus* genotypes, no significant differences in alpha diversity were observed, however, beta diversity analysis revealed a clear separation between wild-type and CSSP mutants in both rhizosphere and root compartments (Fig. S6A,D, Fig. 6C). Additionally, rhizosphere communities of *ccamk* mutants formed a distinct cluster, separate from both wild-type and other mutant genotypes. Several of the most abundant bacterial orders (Fig. S6F), including *Burkholderiales*, *Flavobacteriales*, *Pseudomonadales*, *Xanthomonadales*, and *Rhizobiales*, showed altered relative abundances across *Lotus* genotypes (Fig. 6D, Fig. S6F). In roots, *Rhizobiales* were significantly reduced in CSSP mutants, primarily due to the decline in abundance of symbiotic *Mesorhizobium* isolates, which dominated wild-type roots and nodules but were nearly absent from mutant roots (Fig. 6D,E, Fig. S7A,B). More than 50% of detected isolates showed differential abundance in at least one CSSP mutant relative to wild-type (Fig. 6E, Table S10). These differentially abundant isolates were largely overlapping between rhizosphere and root fractions and spanned all major bacterial orders. Non-symbiotic *Rhizobiales* genera increased in abundance, as well as a range of *Burkholderiales* and *Pseudomonadales*. While some isolates showed consistent changes in abundance across all mutants, others exhibited genotype-specific effects (Fig. 6E).

In *Hordeum*, *ccamk* and *nsp* mutants also exhibited reduced growth (Fig. 6B). Alpha diversity did not differ significantly between compartments or genotypes (Fig. S6B), whereas beta diversity analysis revealed that wild-type assembled communities distinct from those of CSSP mutants, particularly in *ccamk* and *nsp* mutants (Fig. 6C, Fig. S6E). Among the most abundant bacterial orders (Fig. S6G), *Burkholderiales*, *Pseudomonadales*, *Rhizobiales*, *Sphingobacteriales*, and several other orders were differently represented when the CSSP was impaired (Fig. 6D). At the isolate level, a substantial proportion of the detected community (over 40%), spanning a diverse taxonomic range, exhibited genotype-dependent colonisation patterns (Fig. 6E, Table S11), highlighting both shared and host-specific responses.

In summary, the observed effects of impaired symbiosis signalling on plant performance, root exudation, and root-associated microbiota were recapitulated in reconstitution experiments using defined complex SynComs. These results confirm that CSSP genes influence not only symbiotic interactions but also the assembly of commensal root microbiota, with pronounced effects on plant growth. Although members of *Rhizobiales* and *Burkholderiales* were most strongly affected in both hosts, the specific bacterial taxa responding to CSSP impairment differed across mutants and plant species, underscoring broader and host-specific roles of symbiosis signalling pathways in structuring root-associated bacterial communities.

## Discussion

Plants integrate environmental and genetic cues to shape root-associated microbiomes, yet the mechanisms linking host signalling pathways to microbial recruitment remain poorly understood. Here, we show that the CSSP regulates root microbiota composition, plant growth, and exudate profiles in both the legume *Lotus japonicus* and the cereal *Hordeum vulgare*. Soil- and SynCom-grown plants reveal that host species and genotype shape microbial communities, with CSSP-dependent changes in root exudates driving key shifts in microbiome assembly. These findings highlight a central role for symbiosis signalling beyond classical symbioses in shaping plant-microbe interactions.

Long-term soil fertilisation leaves persistent imprints on microbial communities, known as soil microbial memory or legacy effects (74, 75). NPK, PK, and UF regimes generated distinct soil microbiota, with strong beta-diversity shifts while alpha diversity remained largely stable. This indicates that nutrient inputs reorganise community composition without changing overall bacterial richness, likely reflecting functional redundancy and niche construction within the soil microbiome (76–79). Microbiota composition proved to be a robust predictor of plant nutrient status, with predictive ASVs differing between *Lotus* and *Hordeum*, highlighting species-specific recruitment under contrasting soils. These results support the emerging concept that plant-associated microbiomes can provide diagnostic and predictive information on host nutritional state (54, 80–82). While extending this approach to other soils will require testing across different environments and microbial communities, our findings demonstrate the power of integrating soil legacy with host-microbiome interactions to monitor and potentially guide plant nutrition and precision nutrient management.

Beyond soil-driven effects, our results reveal that the CSSP exerts broad, host-specific influence over root-associated microbial communities, extending well beyond its canonical roles in rhizobial and mycorrhizal interactions. Host-specific microbial assembly represents a first layer of community differentiation, as previously established. On top of this, we uncover a second, CSSP-dependent layer, in which plants actively shape microbiota composition, likely through modulation of root exudates. In *Lotus*, disruption of nodule formation and rhizobial communication allowed commensal enrichment and niche replacement. In *Hordeum*, CSSP mutants exhibited extensive microbial shifts despite the absence of AMS, demonstrating conserved effects across hosts. Prior work in legumes documented similar community shifts upon SNF disruption (55–57), and rhizobial Nod-factor production has been linked to microbial connectivity (54). SynCom experiments confirmed these effects are plant-intrinsic, with CSSP genes modulating bacterial recruitment in a gene-specific manner, likely via altered signalling and nutrient allocation. Root exudate profiling revealed extensive metabolic reprogramming in CSSP mutants, with diversity patterns closely mirroring microbial shifts, indicating strong linkage between metabolic outputs and microbial assembly. Conserved effects on phenylpropanoid metabolism (cinnamic acids, flavonoids, coumarins) were observed in both hosts, consistent with mechanistic links between CCaMK-dependent Ca^2+^ signalling and secondary metabolite production (42). Recent work identified an NSP2-MYB transcriptional module that directly activates flavonoid biosynthesis during nodulation, providing mechanistic evidence that symbiosis signalling pathways actively regulate specialised metabolism to shape microbial interactions (83). At the same time, host-specific changes were evident. *Lotus* mutants showed reduced isoflavonoids, aligning with their established role in rhizobial signalling (23), whereas *Hordeum* mutants displayed differential exudation of gibberellin and abscisic acid catabolites, known modulators of AMS signalling, phosphate-dependent symbiosis suppression, and stress responses (84–89). Together, these findings support a model in which CSSP genes coordinate metabolic flux through signalling and defence pathways, fine-tuning chemical outputs that govern microbial recruitment and activity. Divergent metabolic wiring in *Lotus* and *Hordeum* underscores that CSSP regulation of metabolism is host-specific and shaped by evolutionary adaptation.

Among CSSP components, CCaMK emerged as a central hub linking symbiosis signalling to broader microbial regulation. CCaMK impairment led to pronounced shifts in root-associated bacterial communities, forming genotype-specific clusters and altering the abundance of specific ASVs. In *Hordeum*, *ccamk* mutants showed reduced growth in soil and SynCom conditions but grew comparably to wild-type plants under axenic conditions, indicating that CCaMK mediates plant responses to microbial colonisation rather than intrinsic growth. Similar microbe-dependent growth penalties have been observed in AM-defective maize under field conditions (90). These results suggest that CCaMK integrates Ca^2+^ signalling with downstream metabolic outputs to modulate microbial community assembly, balancing colonisation by beneficial microbes while constraining maladaptive interactions. Emerging evidence also links CCaMK to immunity and stress responses, further positioning it as a central regulator of plant-microbe co-regulation (42, 91).

Collectively, our findings demonstrate that CSSP genes orchestrate plant growth, root exudation, and microbial community assembly even under complex soil conditions. Beyond mutant studies, natural variation in CSSP genes offers a promising avenue to exploit plant-microbe interactions for agricultural benefit. By understanding how host genetics shape microbial recruitment and nutrient acquisition, it may be possible to select or engineer crops that efficiently harness beneficial microbiomes, even in non-symbiotic contexts. These results highlight that ancient symbiotic signalling pathways modulate broader microbial networks, offering opportunities to improve plant performance, nutrient use efficiency, and resilience through microbiome-informed breeding and management strategies.

## Materials and Methods

### Plant material, seed sterilisation and germination

*Lotus japonicus* ecotype Gifu-B129 and *Hordeum vulgare* var. Golden Promise were used as wild-type genotypes. Common Symbiosis Signalling Pathway (CSSP) mutants *symrk-3*, *ccamk-13*, *nsp1-1*, and *nsp2-3* were used for *Lotus* (Aarhus University LORE1 insertion mutant collection in the Gifu-B129 background (92)), while CRISPR/Cas9 mutants *symrk-1*, *ccamk-1*, *nsp1-1*, and *nsp2-1* were used for *Hordeum* (49). Seeds were surface-sterilised in 4% (v/v) bleach for 5min, followed by five washes with sterile Milli-Q water. *Lotus* seeds were additionally scarified with sandpaper prior to sterilisation and incubated overnight at 4°C with gentle shaking in sterile Milli-Q. Seeds were germinated on moist filter paper in sterile square Petri dishes for five days (21°C; 14/10h day/night cycle).

### Askov soils

The soils used in this study were sampled from the Askov long-term field experiment at Askov Experimental Station, Southern Denmark (GPS: 55.466 N, 9.117 E), which has maintained defined fertiliser treatments since 1894 (59). In 2019, soil was collected from the B3 field at a depth of 5-15cm from three plots representing different fertilisation regimes: (1) soil receiving a standard dose of nitrogen, phosphorus, and potassium mineral fertilisation (plot 311, here termed NPK), (2) nitrogen-deficient soil receiving only phosphorus and potassium (plot 321, here termed PK), and (3) unfertilised soil (plot 322, here termed UF). The NPK and UF treatments were initiated in 1894, whereas the PK treatment began in 1935.

The Askov soil is a light sandy loam, consisting of 76% sand, 13% silt, and 11% clay, and is classified as an Alfisol (Typic Hapludalf, USDA Soil Taxonomy). The field follows a four-course crop rotation of winter wheat, silage maize, spring barley, and a grass-clover mixture for cutting. Lime is applied every 4-5 years to maintain a soil pH of 5.5-6.5. The mean annual precipitation and temperature at the site are 953mm and 8.8°C, respectively (1999-2018).

### Plant cultivation in Askov soils with different fertilisation regimes

Germinated seedlings were transplanted into sterile 4x4x9cm pots containing a bottom layer of autoclaved LECA (2-4mm; 120°C for 20min; 50ml), an upper layer of soil (80ml), and a 1cm top layer of autoclaved fine vermiculite. Three soils from the long-term Askov experiment (NPK, PK, and UF; see “Askov Soils” section) were used (59). Six biological replicates were included per condition. Each replicate consisted of three pots with one plant per pot for *Hordeum* (total *N* = 18) and ten plants per pot for *Lotus* (total *N* = 60), except for *Lotus symrk-3* mutants, which contained six plants per pot due to reduced germination rates. Plants were watered with sterile Milli-Q water and cultivated in growth chambers with *Lotus* (16/8 day/night hours, 75/75% room humidity day/night, 22/18°C day/night temperature, 200/0μmol m^-2^ s^-1^ light density day/night) or *Hordeum* settings (16/8 day/night hours, 75/80% room humidity day/night, 18/16°C day/night temperatures, 200/0 μmol m^-2^ s^-1^ light density day/night) for three weeks.

### Sample collection and 16S rRNA amplicon sequencing of plants grown in Askov soils

The soil block containing plant roots was removed from the pots, and loosely adhering soil was manually removed. Shoots and roots were separated, and shoot fresh weight was recorded. Roots from plants belonging to one biological replicate were pooled and washed five times in sterile Milli-Q water by vigorous manual shaking for 30sec. ‘Rhizosphere’ samples were obtained from pellets collected after centrifugation of the first wash solution (15min at 4,000 x *g*). For *Lotus* wild-type plants, nodules were excised using scalpels and collected as separate ‘nodule’ fraction. Roots were blotted dry using sterile filter paper, cut into 1cm pieces, and collected as the ‘root’ fraction. Bulk soil samples were obtained from unplanted pots. *Lotus* root samples were stored in sterile 15ml tubes and *Hordeum* root samples in 15ml Precellys tubes containing a single 1x1cm ceramic bead. Rhizosphere and bulk soil samples were transferred to tubes of the Fast DNA Spin Kit for Soil (MP Biomedicals), snap-frozen in liquid nitrogen, and stored at −70°C until further processing.

*Lotus* root samples were homogenised by vortexing until a fine powder was obtained. *Hordeum* roots were homogenised using a Precellys Homogeniser (Bertin Technologies) for 30sec at 6200rpm, repeated if necessary. A subset of homogenised root material was transferred to Fast DNA Spin Kit for Soil tubes (MP Biomedicals), and DNA was extracted according to the manufacturer’s protocol. DNA concentrations were determined using Quant-iT PicoGreen dsDNA assay kit (Life Technologies) and adjusted to 3.5ng/μl. Library preparation and sequencing were performed separately for *Lotus* and *Hordeum* samples. Primers targeting the V5**-**V7 region of the bacterial 16S rRNA gene (799F, 1192R; sequence information in Table S12) were used in an initial PCR followed by a second PCR using unique barcoded primers with Illumina MiSeq (*Lotus*) or NovaSeq (*Hordeum*) adapters for individual samples. For *Lotus* samples, synthetic spike-in DNA (sequence in Table S12; 10^5^ copies) was included in PCR1, but excluded from downstream analyses. Amplicons were purified, pooled, size-selected by gel electrophoresis, and extracted using the Nucleospin Gel and PCR Clean-up Kit (Macherey-Nagel). Libraries were concentrated using AMPure XP beads (Beckman Coulter), quantified using Qubit dsDNA HS and BR Assay Kit (Thermo Fisher Scientific), and sequenced on Illumina MiSeq (Genewiz) for *Lotus* or NovaSeq (Novogene) platforms for *Hordeum* samples.

### Plant cultivation in gnotobiotic systems supplied with synthetic bacterial communities

Two independent gnotobiotic reconstitution experiments were performed. *Lotus* seedlings were cultivated in a closed system of autoclaved magenta boxes filled with a sterile 5:2 mixture of autoclaved LECA and baked sand (overnight at 200°C), topped with a 1cm layer of autoclaved fine vermiculite. For each genotype, four biological replicates were included, each consisting of 25 seedlings per box (total *N* = 100). Plants were supplied with low-nitrogen, low-phosphorus Long Ashton medium (0.75mM MgSO_4_x7H_2_0, 1mM NaNO_3_, 1mM K_2_SO_4_, 2mM CaCl_2_x2H_2_O, 0.0032mM Na_2_HPO_4_, 5μM MnSO_4_, 0.25μM CuSO_4_x5H_2_O, 0.5μM ZnSO_4_x7H_2_O, 25μM H_3_BO_3_, 0.1μM Na_2_MoO_4_x2H_2_O, 25μM Fe-EDTA, pH 6.8) and cultivated for four weeks at 21°C, 16/8 day/night hours, 100/0 μmol m^-2^ s^-1^ light density day/night. One day after planting, plants were inoculated with the Lj-SPHERE bacterial culture collection (93) (Table S8). Individual cultures were pooled in equal proportions based on OD_600_ measurements, diluted to a final OD_600_ 0.02, and 3ml of inoculum were added per box. For *Hordeum*, seedlings were planted in sterile 4x4x9cm pots containing a sterile 5:2 LECA/sand mixture and a top layer of autoclaved vermiculite. Six biological replicates were included per genotype, each consisting of three pots with one plant per pot (total *N* = 18). Plants were supplied with Long Ashton medium and cultivated for three weeks in growth chambers with 16/8 day/night hours, 75/80% room humidity day/night, 22/18°C day/night temperatures, and 200/0μmol m^-2^ s^-1^ light density day/night settings. One day after planting, a *Hordeum*-derived bacterial culture collection (Plant Molecular Biology Group, Aarhus University, DK, Table S9) (73) was applied as inoculum (final OD_600_=0.01; 3ml per pot). In addition, plants were inoculated with AM fungi on the day of planting by applying 1ml of *Rhizophagus irregularis* spores (400 spores per ml, washed twice with Long Ashton medium; Agro Nutrition) 1cm below the soil surface near the roots.

### *Hordeum* growth test in microbe-free condition

To assess for potential growth defects associated with CSSP mutations in *Hordeum*, a microbe-free control experiment was conducted in parallel to the gnotobiotic setup. All genotypes were cultivated in sterile substrate and supplied with Long Ashton medium containing elevated nitrogen and phosphorus concentrations (2.5mM NaNO3 and 1mM Na2HPO4), without microbial inoculation. Ten biological replicates were included per genotype, each consisting of one pot with one plant. Shoot fresh weight was recorded after three weeks of growth.

### Sample collection and 16S rRNA amplicon sequencing of plants grown in gnotobiotic systems

Details on harvest and sample collection are similar to those described for the soil experiment, with minor modifications. In the gnotobiotic studies, leca and sand were removed manually. For *Lotus*, plants from a single magenta box system were pooled for the collection of ‘rhizosphere’ and ‘root’ samples. Roots were ground using a mortar and pestle under liquid nitrogen, and a subset of the ground root material was transferred to tubes of the Fast DNA Spin Kit for Soil (MP Biomedicals) for DNA extraction. For *Hordeum*, roots from each biological replicate were cut into 1-cm pieces, mixed thoroughly, and a subset was used for downstream processing. 16S rRNA V5-V7 amplicon libraries were prepared as described previously and sequenced using Illumina MiSeq (Genewiz) for *Lotus* samples and NovaSeq (Novogene) for *Hordeum* samples. For *Lotus*, spike-in DNA (sequence info in Table S12, 10^4^ copies) was included in PCR1 but was not used for downstream analysis.

### Pre-processing of amplicon sequencing data

Amplicon reads from *Lotus* and *Hordeum* libraries were processed separately using the QIIME2 pipeline (version 2021.4) (94). Quality control and denoising of demultiplexed reads were performed using the DADA2 plugin, generating representative amplicon sequence variants (ASVs) (95). ASVs were filtered to retain those with at least 10 reads in at least four samples. Taxonomic classification of representative sequences was performed using QIIME2’s naïve-Bayes classifier implemented in the scikit-learn-based q2-feature-classifier (96), with reference to the SILVA database (version 138_99) (97). For data obtained from gnotobiotic systems, pre-processing was likewise performed using QIIME2 and the DADA2 plugin for quality control and demultiplexing (94, 95). No abundance filtering was applied, as defined sets of isolates with known sequence information were used. Taxonomic assignment of detected ASVs was performed using VSEARCH with a 99% identity threshold (98). Reference sequences consisted of 16S rRNA V5-V7 regions extracted from whole-genome sequences of strains included in the inoculum (Tables S8, S9). Isolates with identical V5-V7 regions were grouped together. ASVs not matching any member of the SynCom reference set were removed prior to downstream analysis.

### Data analysis, statistics and data visualization

All analyses were performed in R (version 4.5.2). For *Lotus* soil and SynCom datasets, spike-in DNA was removed from the ASV table prior to analysis. For alpha-diversity analyses, samples with fewer than 1000 reads were excluded, and read counts were rarefied to the minimum sequencing depth using phyloseq (version 1.52.0) (99). Alpha diversity was quantified using Chao1 index as computed in QIIME2. For beta-diversity analyses, principal coordinate analysis (PCoA) (100) based on Bray-Curtis dissimilarities (101) was performed separately for each genotype. Constrained analysis of principal coordinates (CPCoA (102); capscale function in the vegan package in R (103)) using Bray-Curtis dissimiliarities was conducted with genotype and soil type as constraining variables. Permutation tests were used to assess the effects of the constrained variables on bacterial community composition. Relative ASV abundances were calculated using total sum scaling (TSS). Differential abundance analysis of individual ASVs was performed using MaAsLin2 (104) and a structural-zero filtering step as proposed by Kaul *et al*. (105), with the additional requirement that the non-structural-zero group contained at least 20 reads across at least three samples. P-values were adjusted for multiple testing using the Benjamini-Hochberg procedure (106), with a false discovery rate (FDR) of 5%. Pairwise significance testing was performed using ANOVA and Tukey post-hoc test. Data visualization was carried out in R using ggplot2 (107). A full list of used packages and versions can be found in Table S13.

### Nutrient prediction analysis of plants cultivated in Askov soils

Prediction of soil type was performed using Compositional Data via Continuos Relaxations (codacore) (108). Because codacore supports only binary categorical variables, a nested modelling scheme was applied to predict the trinary soil type variable (NPK, PK, UF). Two models were fitted: one distinguishing NPK from non-NPK soils, and a second distinguishing PK from UF soils. The first model was trained on data from all three soils, whereas the second was trained only on PK and UF samples. As illustrated in Figure 3A, the first model predicts whether a sample originates from NPK soil; if not, the second model predicts whether it originates from PK or UF soil. Two-thirds of the data were used for model training and one-third was used to assess prediction accuracy. The split was performed randomly while ensuring equal representation of all three soils and all five genotypes in both training and test sets. Separate nested models were trained for *Lotus* and *Hordeum* datasets. Analysis only included bacterial ASVs.

### Root exudate experimental setups

For *Lotus*, sterile-germinated wild-type and CSSP mutant seedlings were transferred to sterile square Petri dishes half-filled with solid quarter-strength B&D medium without nitrogen (1.4% Agar Noble, Difco) (109). For each genotype, six biological replicates were prepared, with 12 plants per plate. A sterile filter paper was placed between the roots and the medium, and plants were fixed using an autoclaved aluminium foil grid. Plates were maintained vertically in custom plastic racks covering the lower half of the plates to keep roots in darkness. Plants were cultivated at 21°C, 16/8 day/night hours, 100/0 μmol m^-2^ s^-1^ white light day/night for two weeks.

For *Hordeum*, sterile-germinated wild-type and CSSP mutant seedlings were cultivated in autoclaved Duran borosilicate glass tubes containing 50ml sterile VWR sand (CAS 14808-60-7, washed with sterile Milli-Q and baked for 4h at 200°C), and supplemented with 15ml of low-nitrogen, low-phosphorus Long Ashton medium (as used for the *Hordeum* gnotobiotic experiment). One seedling was added per tube, which was closed with an autoclaved foam plug. Unplanted sand-only controls were included. Plants were cultivated under the same conditions as *Lotus* for three weeks.

### Root exudate harvest, sample preparation and LC-MS/MS

For *Lotus*, plants from a single Petri dish were pooled and transferred to a sterile glass vial containing 20ml sterile Milli-Q. After 3h, plants were removed, and the solution was filter-sterilised using 0.4µm cell strainers. For *Hordeum*, plants were gently removed with sterile tweezers. Thirty millilitres of sterile Milli-Q were added to the sand, tubes were sealed with Parafilm, vortexed for 30s, and the suspension was allowed to settle. The supernatant was collected into fresh tubes and filtered through 0.4µm cell strainers. All samples were stored at −70°C and subsequently freeze-dried in a lyophiliser (CD52-I, HETOSIC, Heto Lab Equipment) for 1-2 days. For *Lotus*, freeze-dried samples were spiked with 40µl internal standard (10.6µg/ml vanillin-(phenyl-13C6) in methanol), reconstituted in 5ml methanol, shaken at 1500rpm for 5min, and sonicated for 10min at room temperature. Extracts were filtered through hydrophobic PTFE filters (0.22µm pore size, 25mm), dried under a gentle nitrogen flow, up-concentrated with 200µl D6-abscisic acid in 80% methanol, and transferred to vials with 300µl inserts. For *Hordeum*, freeze-dried samples were spiked with 20μl 10μg/ml ketopinic acid in methanol and dried under nitrogen flow. Samples were reconstituted in 3ml methanol, sonicated for 10min, and filtered using hydrophobic PTFE syringe filters. After drying under nitrogen, samples were reconstituted in 300μl of 80:20 (v/v) methanol:Milli-Q water containing 1μg/ml camphoric acid (internal standard) and transferred to vials for analysis. Analyses for both species were performed using ultra-high-performance liquid chromatography (UHPLC; Dionex Ultimate 3000, Thermo Fisher) coupled to a quadrupole time-of-flight mass spectrometer (qToF-MS; Bruker Compact) with electrospray ionisation in negative mode. Separation was achieved using an ACQUITY UHPLC HSS T3 column (2.1x1000 mm, 1.8μm; Waters) at a flow rate of 0.2ml/min. Mobile phases consisted of 0.1% acetic acid in Milli-Q water and 0.1% acetic acid in acetonitrile, and a linear gradient program, as described by Salomonsen *et al*., 2024 (110), was applied. A 2μl injection volume was used, and the column temperature was maintained at 45°C.

### Metabolite data analysis, statistics and visualization

Raw LC-MS data were pre-processed using MzMine 3.9.0 (111). Features with retention times greater than 19min were excluded. Chemical annotation was performed using CANOPUS implemented in SIRIUS 6.3.4, as well as the ClassyFire and NPClassifier tools (112–115). Prior to statistical analyses, metabolites detected in fewer than 10% of samples were removed. Differential exudation between sand control samples and the five genotypes (wild-type, *symrk*, *ccamk*, *nsp1*, and *nsp2*) was assessed using Tobit regression (116), with the minimum non-zero measurement used as the lower detection limit, followed by a likelihood-ratio test with Benjamini-Hochberg correction at an FDR of 5%. For metabolites detected in all samples, this approach is equivalent to linear regression. Principal component analysis (PCA) (117) was subsequently performed. Differential exudation between wild-type and each mutant was analysed using the same Tobit regression framework, likelihood-ratio testing, and BH correction (FDR 5%). Metabolites with annotation probabilities below 60% were classified as unknown.

### Data and code availability

All data and code will be made publicly available upon publication of the final peer-reviewed manuscript.

## Acknowledgements

We thank Dr. Sha Zhang, Dr. Florian Lamouche, and Dr. Simon Kelly for valuable input on experimental design and data analysis, and Dr. Eleni Soumpourou for assistance with the import and genotyping of *Hordeum* mutant seeds. This work was supported by grants made to the University of Cambridge by Gates Foundation and the UK Foreign, Commonwealth and Development Office (OPP1028264) and Gates Agricultural Innovations (INV-57461) known as the Enabling Nutrient Symbioses in Agriculture (ENSA) project, the Novo Nordisk Foundation as part of the “Molecular Mechanisms and Dynamics of Plant-microbe interactions at the Root-Soil Interface” project (NNF19SA0059362), and by the Carlsberg Foundation and the Danish National Research Foundation (DNRF 172) through the Center of Excellence for Chemistry of Clouds. Figures were created in part with BioRender.com.

## Author Contributions

A.M., I.T.J, C.L.S, M.G. and S.R. designed the experiments. A.M. performed the experiments with technical support from Z.B., K.T. and H.D. LC-MS/MS was performed by C.L.S, Þ.N.B. and M.G. Data analysis was performed by A.M., I.T.J. and C.L.S. A.M. wrote the first manuscript draft with following input from all authors. *Hordeum* symbiosis mutants were provided by G.O. Askov soils were provided by B.T.C.

## Supplementary Information

### Supplementary Figures

**Figure S1.**
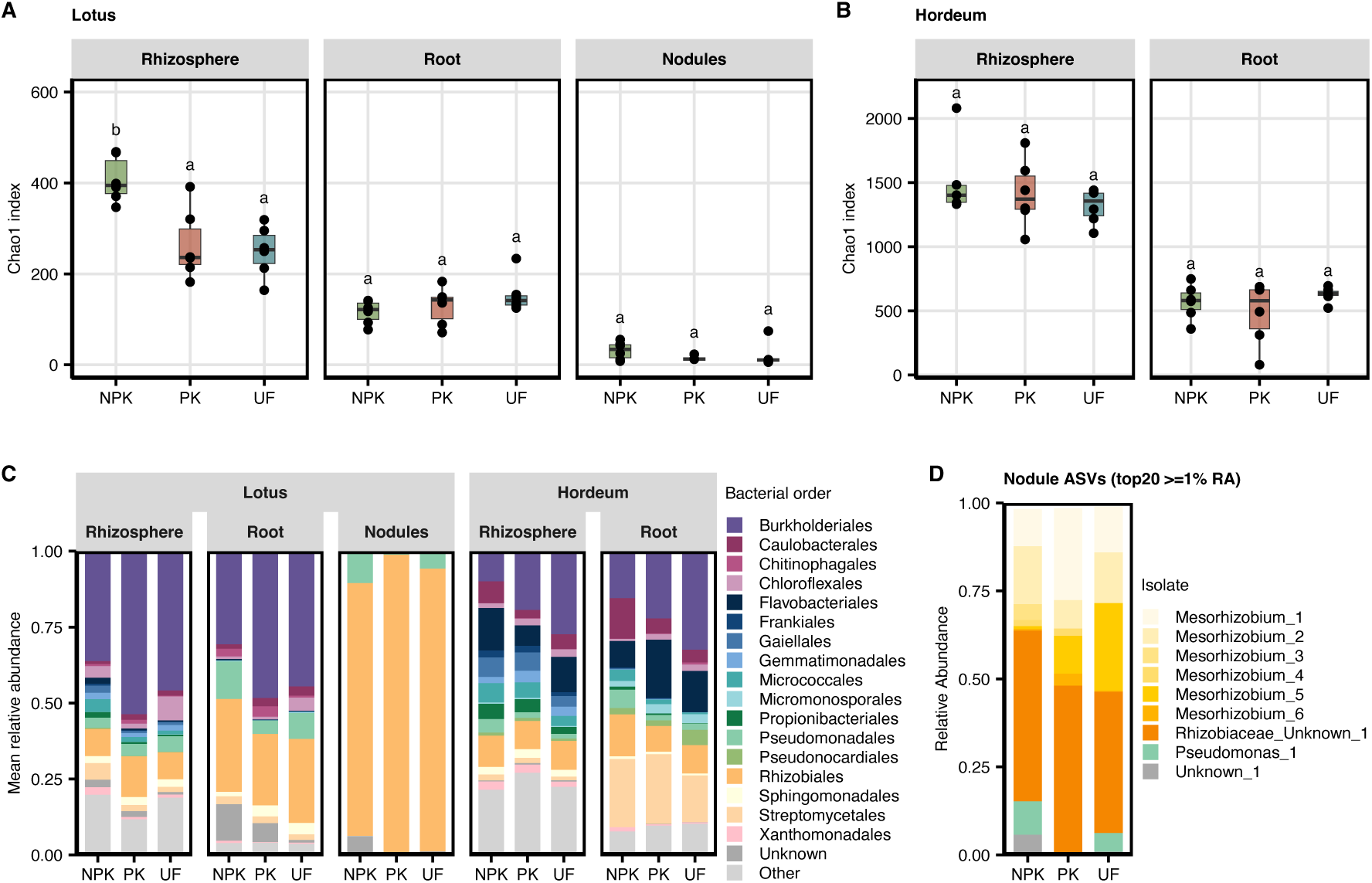
Microbial diversity and composition in rhizosphere, root, and nodule compartments of *Lotus japonicus* and *Hordeum vulgare*. Samples were collected as described in Figure 2. (A,B) Chao1 alpha diversity of bacterial communities in *Lotus* (rhizosphere, root, and nodule) and *Hordeum* (rhizosphere and root) compartments across three soil types (NPK, PK, UF). Letters indicate significant differences between soils within each compartment (one-way ANOVA followed by Tukey’s HSD, P < 0.05). (C) Stacked barplots showing mean relative abundances (RA) of bacterial orders with mean RA ≥1% across all rhizosphere and root samples in *Lotus* or *Hordeum*. Lower-abundance orders are grouped as “Other”. (D) Stacked barplots of *Lotus* nodule ASVs. The most abundant ASVs per soil with ≥1% mean relative abundance were plotted; less abundant ASVs were excluded. Unknown genera were annotated based on family information. Colours correspond to genera/ASVs.

**Figure S2.**
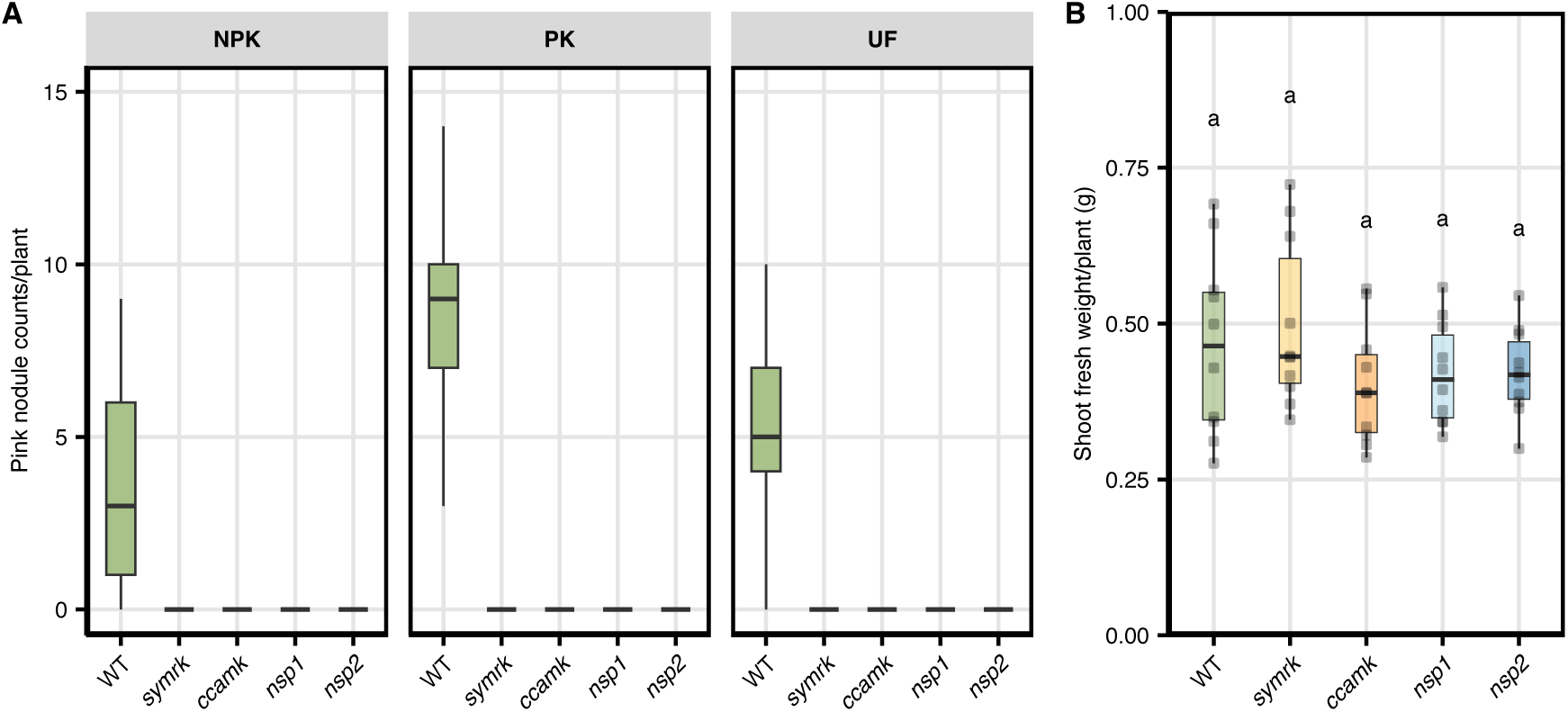
Nodule counts in *Lotus* genotypes and microbe-free growth test of *Hordeum* CSSP mutants. (A) Pink nodule counts per *Lotus* plant grown for three weeks in NPK, PK, or UF soils. Data are shown for wild-type and symbiosis mutants, though mutants did not form nodules. Sample collection and experimental setup followed the procedure described in Figure 4. (B) Shoot fresh weight (g) for *Hordeum* wild-type and symbiosis mutants (*N* = 10, one plant per pot per genotype) grown under sterile, microbe-free conditions with elevated nitrogen and phosphorus (2.5mM NaNO_3_, 1mM Na_2_HPO_4_). Statistical significance was assessed using one-way ANOVA followed by Tukey’s HSD (P < 0.05).

**Figure S3.**
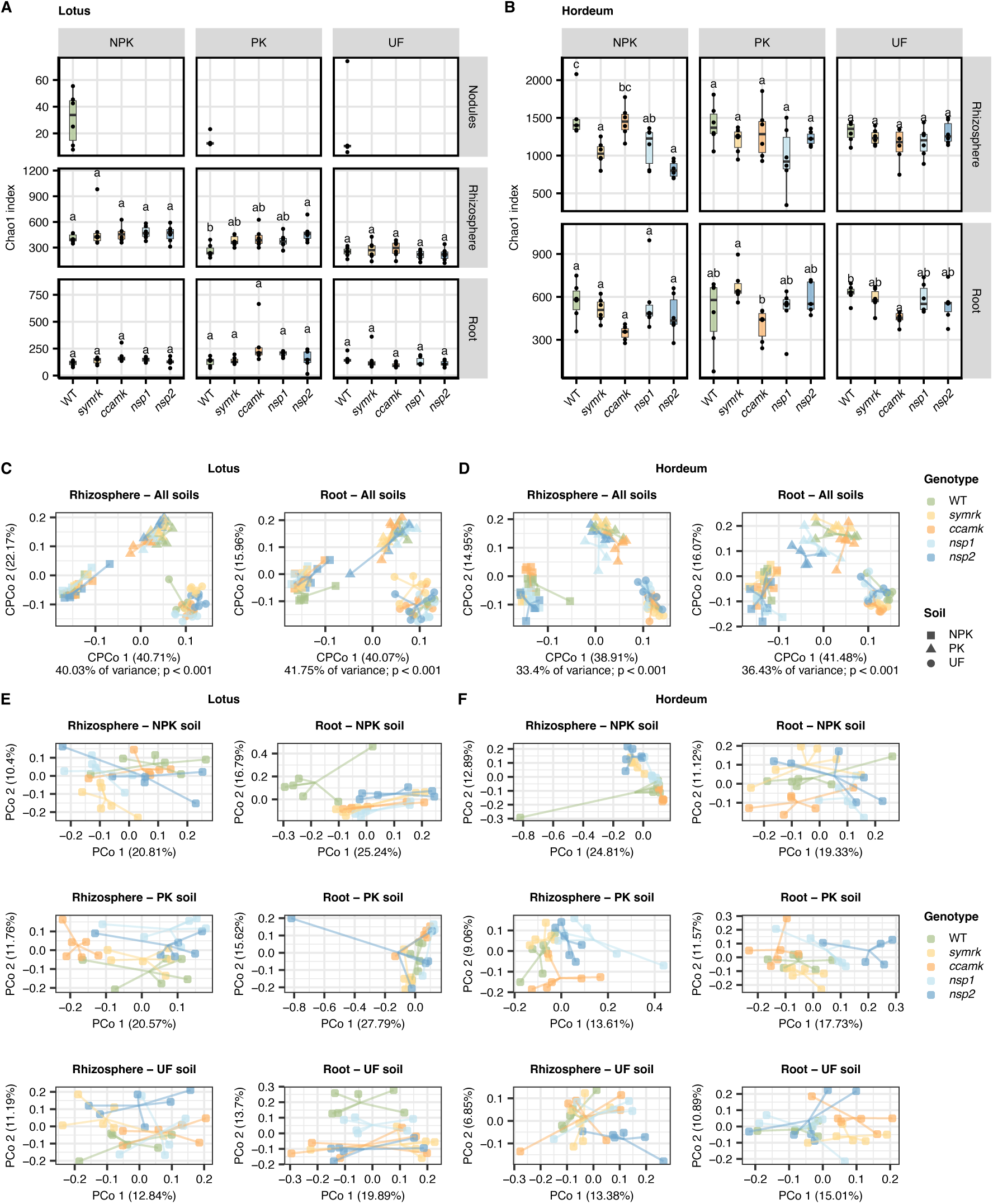
Microbial diversity and composition in *Lotus* and *Hordeum* wild-type and CSSP mutants across soils. Experimental setup is described in Fig. 4. (A) Chao1 alpha diversity of bacterial communities in *Lotus* rhizosphere, root, and nodule compartments across NPK, PK, and UF soils for all genotypes. (B) Chao1 alpha diversity in *Hordeum* rhizosphere and root compartments across soils for all genotypes. Statistical significance across genotypes within each compartment-soil combination (A,B) was assessed using one-way ANOVA followed by Tukey’s HSD (P < 0.05). (C,D) Constrained PCoA of Bray-Curtis distances showing bacterial community composition in rhizosphere and root compartments across all soils, with genotypes indicated by colours and soil types by shapes (C: *Lotus*; D: *Hordeum*). (E,F) Unconstrained PCoA of Bray-Curtis distances for each compartment-soil combination, with colours indicating genotypes (E: *Lotus*; F: *Hordeum*).

**Figure S4.**
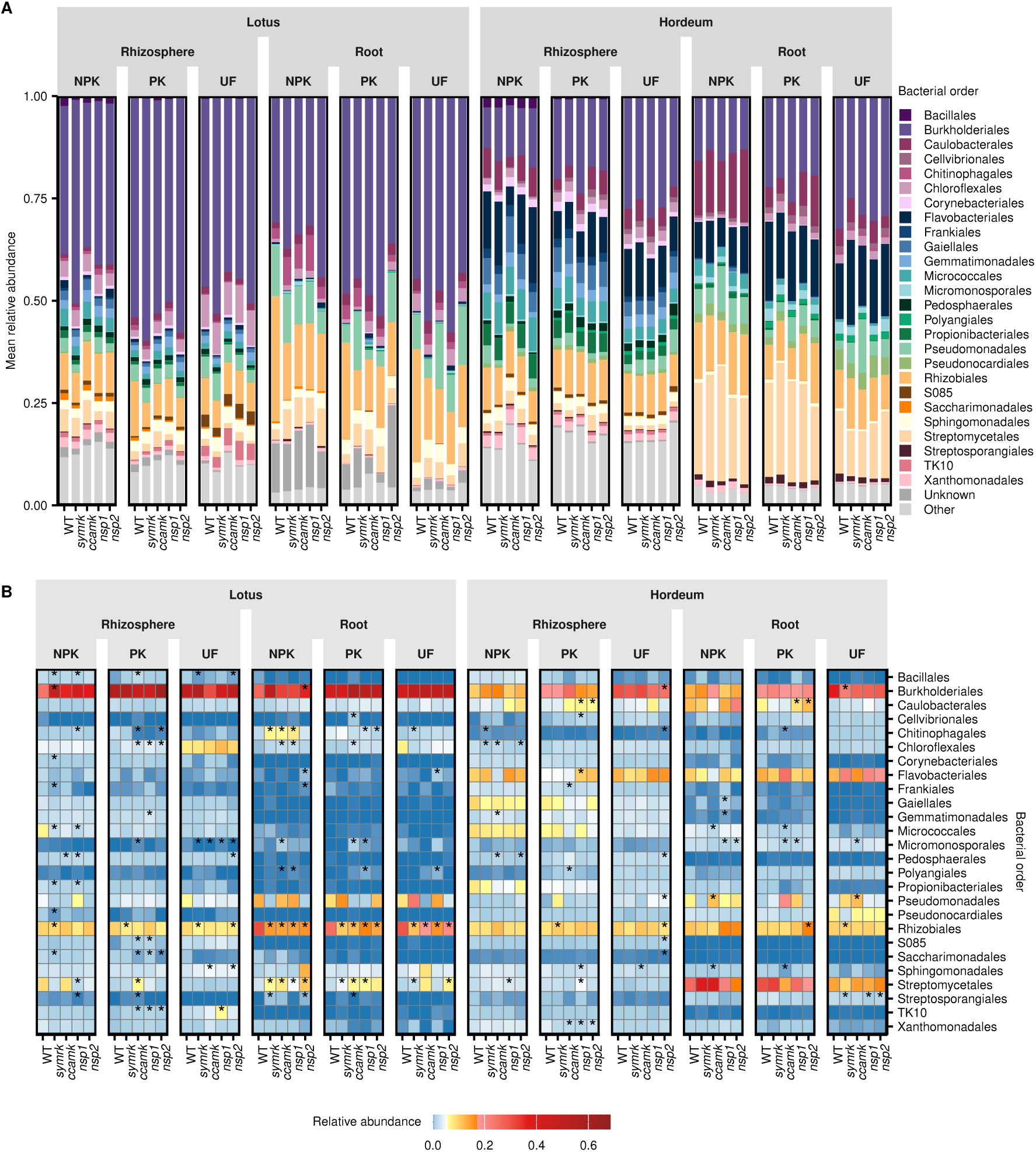
Relative abundance of dominant bacterial orders in *Lotus* and *Hordeum* wild-type and CSSP mutants across soils. Experimental setup is described in Fig. 4. (A) Stacked barplots showing mean relative abundance of bacterial orders across plant-compartment-soil-genotype combinations. The top 20 bacterial orders (based on mean relative abundance in wild-type samples of *Lotus* and *Hordeum*) are shown, with remaining orders grouped as “Other”. (B) Heatmap showing mean relative abundance of bacterial orders across the same combinations, with colours ranging from low abundance (dark blue) to high abundance (red). Asterisks indicate orders with significantly different relative abundance compared with the wild-type within each plant-compartment-soil combination, assessed using linear regression (adjusted P < 0.05).

**Figure S5.**
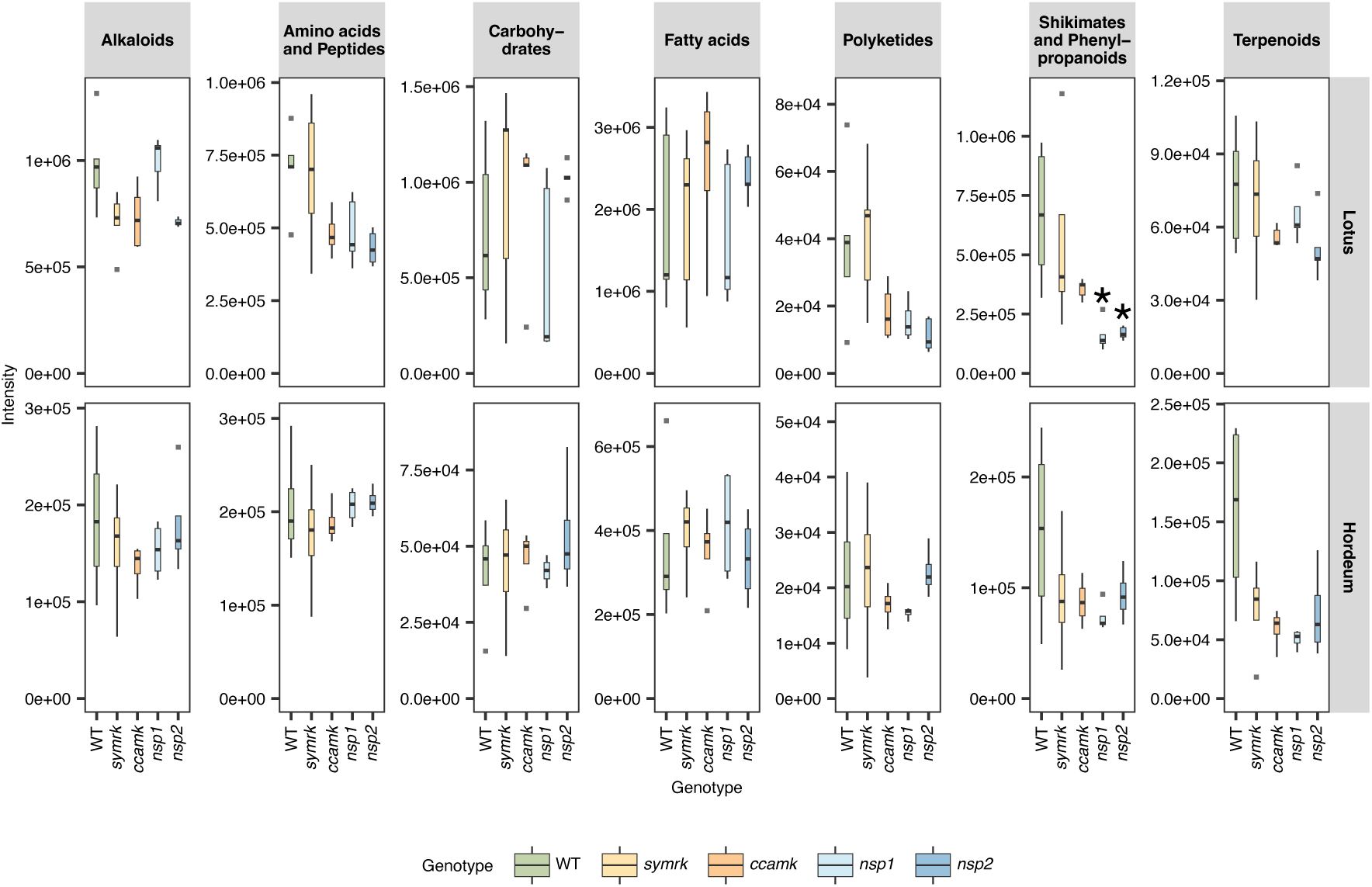
Cumulative pathway-level intensities of root exudates in *Lotus* and *Hordeum* wild-type and CSSP mutants. Experimental setup is described in Figure 5. Boxplots show the cumulative intensity of metabolites grouped by predicted pathway (NPC classifier) for each genotype and host species. An asterisk indicates a statistically significant difference compared to wild-type (linear regression with t-tests, P < 0.05 after correction for multiple testing with Benjamini Hochberg).

**Figure S6.**
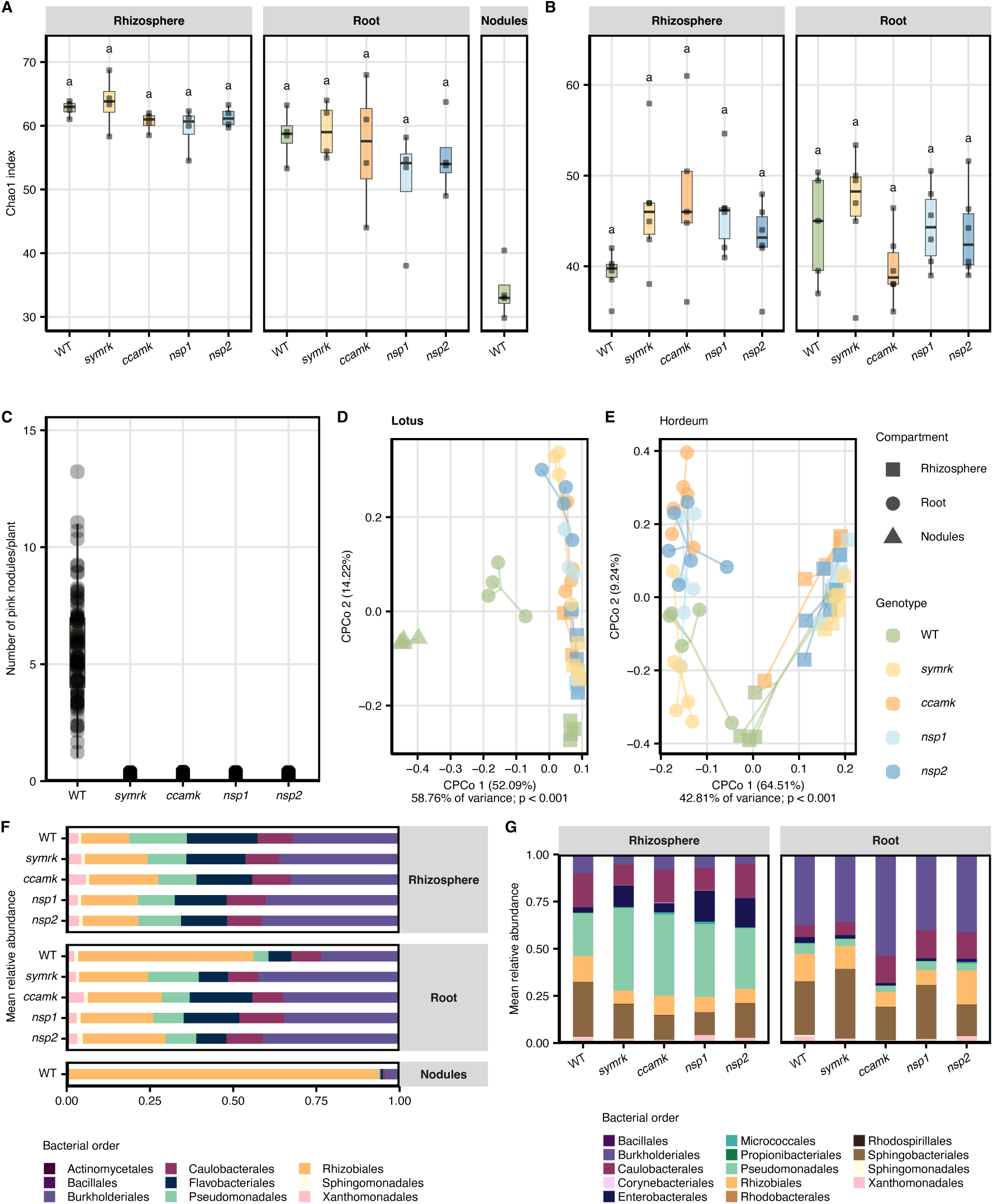
Diversity and compositional analysis of bacterial communities across compartments and genotypes in SynCom experiments. Experimental setup is as described in Fig. 6. (A,B) Chao1 alpha diversity of bacterial communities in (A) *Lotus* (rhizosphere, root, and nodule) and (B) *Hordeum* (rhizosphere and root) compartments across wild-type and symbiosis mutants in the SynCom experiments. (C) Pink nodule counts in *Lotus* wild-type and symbiosis mutants. No nodules were observed in the mutants. (D,E) Constrained PCoA based on Bray-Curtis distances, showing bacterial community composition across genotypes (colours) and compartments (shapes) (D: *Lotus*; E: *Hordeum*). (F,G) Stacked barplots showing mean relative abundances of bacterial orders in each compartment-genotype combination for (F) *Lotus* and (G) *Hordeum*. Unmatched ASVs were excluded from the analysis.

**Figure S7.**
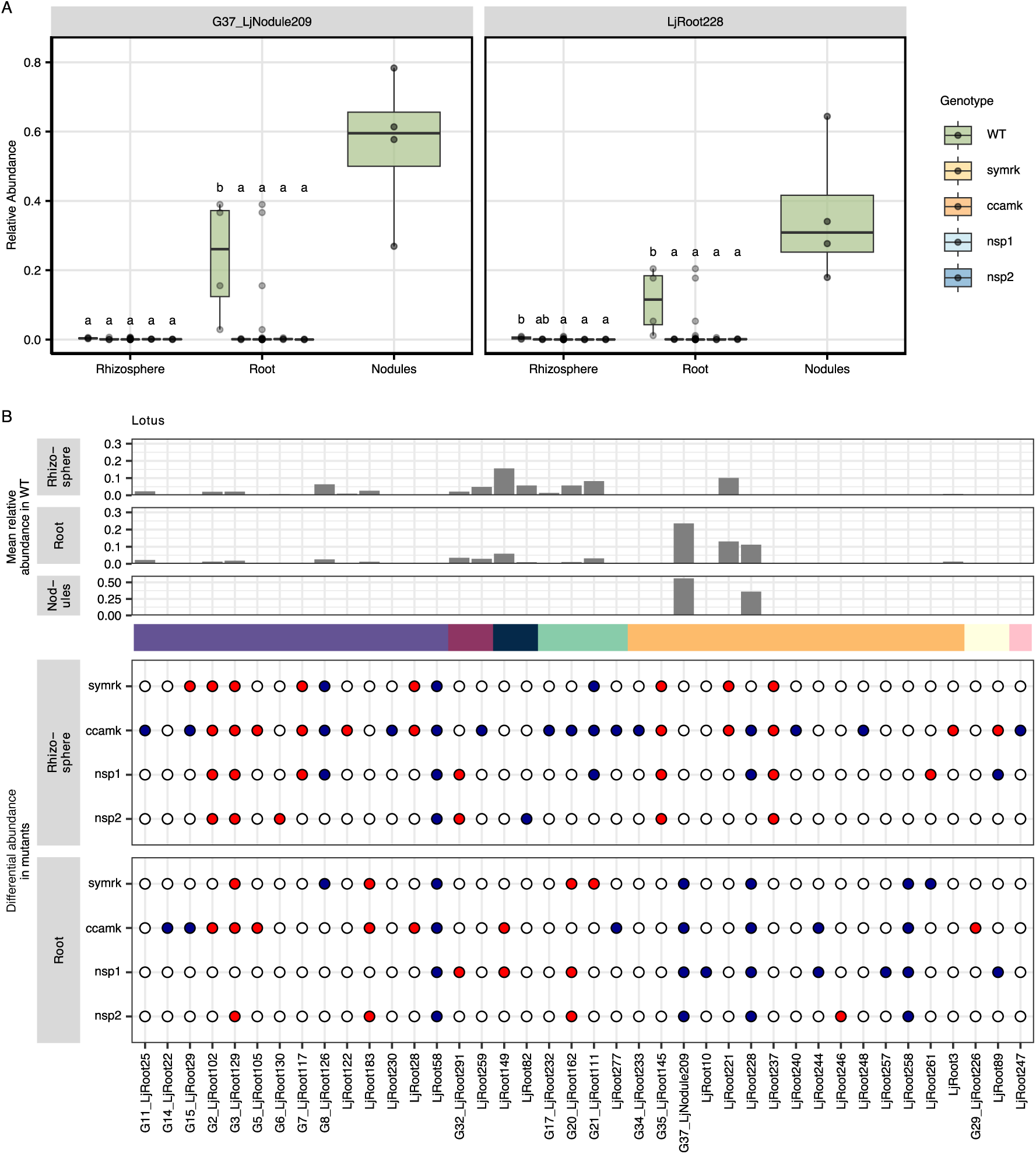
Differential colonisation of commensal and symbiotic bacteria across *Lotus* genotypes in the SynCom experiment. Experimental setup is as described in Fig. 6. (A) Boxplots showing relative abundance differences of bacterial symbionts between *Lotus* wild-type and symbiosis mutants across rhizosphere, root, and nodule compartments. Symbionts include isolate Group37, representing LjNodule209 together with isolates sharing an identical 16S V5-V7 region (Table S8), and isolate LjRoot228. Significance was assessed using one-way ANOVA followed by Tukey’s HSD (P < 0.05). (B) Differentially abundant isolates detected in at least one mutant relative to wild-type based on MaAsLin2 and structural zero analysis. Top panels show mean relative abundance in wild-type rhizosphere, root, and nodule samples. The annotation bar indicates taxonomic assignment at the order level. Bubble plots indicate significant depletion (blue) or enrichment (red) of isolates in mutants relative to wild-type. This plot is identical to Fig. 6E, with the nodule compartment included. Unmatched ASVs were excluded from the analysis.

### Supplementary Tables

**Table S1. Overlap of ASVs across soil types.** ASVs were identified from bulk soil samples (*N* = 3 per soil) as described in Figure 1. An ASV was considered present in a soil if its relative abundance was >0 in at least 2 out of 3 samples. The table lists ASV identifiers, taxonomy information, confidence of taxonomic annotation, and category of occurrence across soils. Categories indicate whether an ASV was detected only in one soil (NPK only, PK only, UF only), two soils (NPK & PK, NPK & UF, PK & UF), or all soils (NPK, PK, UF).

**Table S2. ASVs contributing to soil type prediction models.** ASVs selected by codacore as log-ratio components for the nested soil type prediction models shown in Figure 3. The table lists ASV identifiers, their role in the log-ratio (numerator or denominator), prediction task (NPK *vs*. non-NPK or PK *vs*. UF), plant compartment, and host species. Mean relative abundance (RA) across soils (NPK, PK, UF) is shown, along with taxonomic annotation and confidence scores for taxonomic assignment.

**Table S3. Differentially abundant (DA) ASVs in *Lotus* and *Hordeum* wild-type and CSSP mutants across soils.** The table includes four sheets. Sheet 1 (“Lotus DA overview”) lists, for each *Lotus* genotype-compartment-soil combination, the total number of ASVs detected and the number of DA ASVs relative to wild-type. Sheet 2 (“Lotus DA ASVs”) provides ASV-level information, including ASV identifier, soil, compartment, genotype, log fold change relative to wild-type, method of detection (DA_by_sig from MaAsLin2 or DA_by_struc_zero from structural zero analysis), and taxonomic information. Sheets 3 and 4 contain equivalent overview and ASV-level information for *Hordeum*.

**Table S4. Metabolites detected in *Lotus* root exudates after background filtering.** Feature table of metabolite intensities across *Lotus* samples, including only metabolites enriched in at least one genotype (wild-type, *symrk*, *ccamk*, *nsp1*, or *nsp2*) compared to sand control samples, as assessed using Tobit regression (Benjamini-Hochberg adjusted P < 0.05). The ‘id’ column indicates the metabolite identifier; remaining columns indicate metabolite intensities per sample, named according to their corresponding sample IDs.

**Table S5. Metabolites detected in *Hordeum* root exudates after background filtering.** Feature table of metabolite intensities across *Hordeum* samples, including only metabolites enriched in at least one genotype (wild-type, *symrk*, *ccamk*, *nsp1*, or *nsp2*) compared to sand control samples, as assessed using Tobit regression (Benjamini-Hochberg adjusted P < 0.05). The ‘id’ column indicates the metabolite identifier; remaining columns indicate metabolite intensities per sample, named according to their corresponding sample IDs.

**Table S6. Differentially exuded metabolites (DEMs) in *Lotus* CSSP mutants compared with wild-type.** Table corresponds to the data presented in Fig. 5. Columns include the metabolite FeatureID, the differential exudation status in each mutant relative to wild-type (−1: significantly depleted, 0: no significant change, 1: significantly enriched), and measured intensities in individual samples for all genotypes. Full annotation data are provided for two classifiers (NPC and ClassyFire), along with spectral information (ion mass, retention time, formula, aligned feature ID, etc.). Statistical significance was determined using Tobit regression with correction for multiple testing using Benjamini-Hochberg.

**Table S7. Differentially exuded metabolites (DEMs) in *Hordeum* CSSP mutants compared with wild-type.** Table corresponds to the data presented in Fig. 5. Columns include the metabolite FeatureID, the differential exudation status in each mutant relative to wild-type (−1: significantly depleted, 0: no significant change, 1: significantly enriched), and measured intensities in individual samples for all genotypes. Full annotation data are provided for two classifiers (NPC and ClassyFire), along with spectral information (ion mass, retention time, formula, aligned feature ID, etc.). Statistical significance was determined using Tobit regression with correction for multiple testing using Benjamini-Hochberg.

**Table S8. Lj-SPHERE bacterial culture collection.** The table provides information on isolates from the *Lotus japonicus*-derived bacterial culture collection (Lj-SPHERE) described by Wippel *et al*. (116), which were used in the gnotobiotic *Lotus* experiment (Fig. 6). Sheet 1 lists isolate metadata, including PMS (stock) ID at Aarhus University, isolate ID, collection name, taxonomic annotation from Kingdom to Genus, host species, plant compartment from which the isolate was obtained, and the soil in which the plant was grown. Sheet 2 provides 16S rRNA V5-V7 sequences for the isolates; isolates with identical V5-V7 region were grouped. Sheet 3 lists the isolate group assignments corresponding to these sequence-based clusters.

**Table S9. Hordeum-derived bacterial culture collection.** The table provides information on bacterial isolates obtained from *Hordeum vulgare* roots, used in the gnotobiotic *Hordeum* experiment (Fig. 6). Sheet 1 lists isolate metadata, including PMS (stock) ID at Aarhus University, strain ID, taxonomic annotation from Kingdom to Organism, host species, genome availability, and cultivation medium. Sheet 2 provides 16S rRNA V5-V7 sequences for the isolates.

**Table S10. SynCom isolates with differential abundance in *Lotus* symbiosis mutants.** The table provides an overview of all isolates detected in the *Lotus* SynCom experiment (unmatched ASVs excluded). Separate sheets are included for the rhizosphere and root compartments. For each isolate, the table reports the isolate ID, log fold change in each symbiosis mutant relative to wild-type, and differential abundance status (1 = enriched, −1 = depleted, 0 = not affected). Differential abundance analysis was not performed for isolates present in less than 10% of samples, therefore log fold change and differential abundance status are blank for those isolates. Taxonomic annotation from kingdom to genus level is included, along with relative abundance of each isolate in individual samples.

**Table S11. SynCom isolates with differential abundance in *Hordeum* symbiosis mutants.** The table provides an overview of all detected isolates in the *Hordeum* SynCom experiment (unmatched ASVs excluded). Separate sheets are included for the rhizosphere and root compartments. For each isolate, the table reports the isolate ID, log fold change in each symbiosis mutant relative to wild-type, and differential abundance status (1 = enriched, −1 = depleted, 0 = not affected). Differential abundance analysis was not performed for isolates present in less than 10% of samples, therefore log fold change and differential abundance status are blank for those isolates. Taxonomic annotation from kingdom to genus level is included, along with relative abundance of each isolate in individual samples.

**Table S12. Primer and spike-in DNA sequences used for 16S V5-V7 amplicon library preparation.**

**Table S13. R packages and versions used for data analysis, manipulation, and visualization.**

